# STAT transcription factors regulate host defences and vacuolar integrity during *Mycobacterium marinum* infection

**DOI:** 10.64898/2026.06.09.729547

**Authors:** Justine Toinon, Nabil Hanna, Ludovica Vanzan, Lyudmil Raykov, Thierry Soldati

**Affiliations:** Department of Biochemistry, Faculty of Science, University of Geneva, 30 quai Ernest-Ansermet, Science II, 1211 Geneva-4, Switzerland; Department of Surgery, Faculty of Medicine, University of Geneva, Rue Gabrielle-Perret-Gentil 4, CH-1211 Geneva 14

**Keywords:** Transcription factor, *Dictyostelium discoideum*, *Mycobacterium marinum*, cell-autonomous defences, membrane damage

## Abstract

Signal transducers and activators of transcription (STAT) are central regulators of cytokine-mediated immunity in metazoans, yet their ancestral functions prior to the emergence of interferon signalling remain poorly understood. *Dictyostelium discoideum* expresses four STAT-like factors that impact the intracellular fate of *Mycobacterium marinum* (*Mm*). Using *dst* knockout strains, we identify DstA and DstB as susceptibility factors, whereas DstC acts as a resistance factor, revealing a regulatory axis controlling the integrity of the *Mm*-containing vacuole (MCV). Despite the absence of canonical cytokine receptors and JAK kinases, DstA rapidly translocates to the nucleus in a damage-dependent manner, whereas DstB and DstC exhibit delayed cytosolic redistribution. DstA associates with VacA, a membrane microdomain component, regulates its transcription and accumulation at the MCV, while VacA acts as a cytoplasmic anchor that limits DstA nuclear translocation. Our findings establish STAT-like proteins as evolutionarily ancient regulators of vacuole integrity and host defence, providing new insights into the origins of STAT-mediated immune responses during mycobacterial infection.

## Introduction

Innate immunity, mediated by professional phagocytes such as neutrophils, macrophages, and dendritic cells, constitutes the first line of defence against invading pathogens. These cells detect conserved microbial patterns and rapidly engulf pathogens, eliciting robust intracellular signalling, that drive extensive transcriptional and functional reprogramming. Such cell-autonomous defence programmes must be tightly coordinated to ensure an effective and balanced immune response throughout the course of infection ^1^. Beyond intracellular responses, pathogen recognition also initiates dynamic intercellular communication networks that orchestrate antimicrobial defence, immune tolerance and pathogen clearance. In mammals, signal transducers and activators of transcription (STATs) are central regulators of these processes ^2^. Upon pathogen sensing, cytokine synthesis and secretion mediate cell-to-cell communication and activate signalling cascades that converge on STAT transcription factors, ultimately promoting coordinated immune activation and efficient pathogen elimination.

The STAT signalling pathway was first identified through a study on interferon (IFN)-induced signal transduction ^3^. Since this discovery, numerous cytokines, including IFNs, interleukins, and colony-stimulating factors, have been shown to bind cell-surface receptors and promote receptor dimerization ^4^. A central component of this pathway is the Janus kinase (JAK) family, which comprises non-receptor tyrosine kinases ^5^. Ligand binding to the receptor induces JAK transphosphorylation, followed by tyrosine phosphorylation of the receptor by the activated kinase. These phosphorylation events create docking site for STAT proteins, enabling their recruitment and subsequent tyrosine phosphorylation by JAK. Phosphorylated STATs dissociate from the receptor and form dimers through SH2-domain-phosphotyrosine interactions. These dimers translocate to the nucleus, where they bind target gene promoters and regulate transcription.

Mammalian STAT proteins are central regulators of the host response to bacterial infection, orchestrating immune programs that can be either host-protective or subverted by the pathogen depending on the STAT isoform activated and the cellular context. Collectively, they promote antimicrobial peptide expression, enhance macrophage bactericidal activity, and fine-tune cytokine balance, tissue repair, and immune tolerance ^6,7^. This functional duality is particularly evident during infection with *Mycobacterium tuberculosis* (*Mtb*), the causative agent of tuberculosis. In this context, STAT1 and STAT4 act as central upstream regulators of interferon-driven signalling, whereas STAT3 serves as a major modulator of both pro- and anti-inflammatory responses ^8–10^. During the early phase of infection, STAT1 promotes the transcription of pro-apoptotic factors, which is thought to facilitate the elimination of infected cells by the innate and adaptive immune systems ^11,12^. At later stages, however, the accumulation of unphosphorylated STAT1 through a STAT1-dependent feedback loop enhances anti-apoptotic gene expression, facilitating bacterial persistence and immune evasion ^13^. In parallel, *Mtb*-induced activation of STAT3 in human macrophages suppresses nitric oxide synthesis, thereby establishing a permissive intracellular niche that supports early intracellular replication ^14^. Notably, STAT3 activation has been linked to the activity of early secreted antigenic target 6-kDa (ESAT-6, hereafter EsxA), a major mycobacterial virulence factor secreted by the ESX-1 type VII secretion system ^15^.

*Mycobacterium marinum* (*Mm*), a close genetic relative of *Mtb*, shares highly conserved virulence determinants, including the ESX-1 secretion system and its effectors, and therefore constitutes a powerful model for investigating mycobacterial pathogenesis ^16,17^. As a natural pathogen of freshwater vertebrates, *Mm* recapitulates key pathogenic features of tuberculosis, most notably the formation of organized granulomas in zebrafish that closely resemble human tuberculous lesions ^18,19^. Across multiple host models, STAT signalling has likewise emerged as a critical determinant of the host response to *Mm* infection. In zebrafish, STAT6-dependent signalling is essential for granuloma formation and, through a macrophage cell-autonomous mechanism, drives epithelioid transformation required for granuloma organization ^20^. Beyond granuloma architecture, STAT signalling also regulates infection-associated tissue responses, revealing an important role in host tissue remodelling and vascular pathology ^21^. In *Drosophila*, JAK/STAT activation controls lipid droplet homeostasis within infected phagocytes, highlighting a broader function in immunometabolic adaptation to mycobacterial infection ^22^.

The social amoeba *Dictyostelium discoideum* (*Dd*) constitutes a powerful host model for studying intracellular infection owing to its evolutionarily conserved phagocytic pathway and cell-autonomous defence mechanisms with mammalian innate immune phagocytes ^23–25^. Accordingly, the *Dd*-*Mm* infection system provides a highly tractable and biologically relevant model for dissecting the cellular and molecular determinants of mycobacterial pathogenesis ^26,27^. Following uptake by phagocytes, *Mm* is initially enclosed within a nascent phagosome that rapidly matures and acidifies, transiently exposing the bacterium to a hostile intravacuolar environment characterized by lysosomal hydrolases, reactive oxygen species (ROS), and toxic metal accumulation, all of which contribute to bacteria killing ^24,25,28^. To evade these antimicrobial defences, *Mm* actively remodels the nascent phagosome into a specialized *Mycobacterium*-containing vacuole (MCV) that diverts from the phagolysosomal pathway to create a permissive intracellular niche. An essential mechanism responsible for this evasion is the induction of damage to the MCV membrane through the action of the ESX-1-secreted virulence factor EsxA ^26,29–31^. This process is crucial for mycobacterial intracellular survival and ultimately results in the escape of *Mm* and *Mtb* to the cytosol, a prerequisite for further bacterial growth and dissemination to neighbouring cells ^26^. In response to membrane disruption, host cells deploy repair pathways mediated by endosomal sorting complexes required for transport (ESCRT) and autophagy machineries to contain MCV membrane damage and preserve vacuolar integrity ^32^. In mammals, membrane ATG8ylation, corresponding to the covalent lipidation of ATG8/LC3-family proteins to membranes, has recently emerged as a broad membrane stress, damage, and remodelling response implicated in antimicrobial defence against *Mtb* ^33,34^. This process engages canonical autophagy, LC3-associated phagocytosis, xenophagy as well as membrane repair ^33–35^. Although non-canonical autophagy processes linked to ATG8ylation have not yet been formally characterized in *Dd*, membrane damage induced during *Mm* infection triggers local ubiquitination, and the recruitment of p62/sqstsm1, Atg18 and Atg8-positive structures ^32,36,37^. Central to this response, Ca^2+^-leakage from damaged MCVs activates PefA, an ALG-2 ortholog ^38^ which in turn recruits the E3 ubiquitin ligase TrafE to coordinate the assembly of both ESCRT and autophagy repair machineries at sites of membrane injury ^36,37^. However, mycobacterial lipidic virulence factors, such as phthiocerol dimycocerosates (PDIMs), promote the progression from repairable to catastrophic membrane damage, thereby overwhelming these repair mechanisms, and resulting in bacteria escape to the cytosol ^39,40^. Cytosolic bacteria are subsequently targeted by a selective autophagy process known as xenophagy ^32,36,37^. Consistent with the central role of EsxA in these processes, an *Mm* mutant lacking the RD1 locus (ΔRD1), and therefore deficient in EsxA secretion, induces markedly reduced damage to the MCV membrane, remains confined within the vacuole, and exhibits severely attenuated intracellular survival ^25,31^. This phenocopies the attenuation observed of analogous *Mtb* mutants as well as of the vaccine strain *M. bovi*s BCG ^31,41^. At the host membrane level, sterol-enriched microdomains containing vacuolins (VacA, VacB, and VacC), flotillin-like proteins that stabilize membrane microdomains, accumulate at the MCV and are required for efficient EsxA-mediated membranolysis, thereby promoting bacteria escape from the vacuolar compartment ^42^.

Although *Dd* lacks identifiable homologues of cytokine receptors and JAK kinases, and does not engage canonical interferon signalling, it encodes four STAT-like transcription factors, named Dst proteins (DstA, DstB, DstC, and DstD). Here, we investigate how Dst-dependent signalling activities shape host responses throughout the course of *Mm* infection and define previously uncharacterized non-canonical functions of these Dst proteins. We show that DstA and DstB function as susceptibility factors, whereas DstC acts as a resistance factor, highlighting functional specialization within this STAT-like family. During infection, Dst proteins undergo dynamic nucleo-cytoplasmic redistribution and coordinate distinct host restriction mechanisms. DstA emerges as a damage-responsive regulator of mycobacterial infection, controlling MCV membrane integrity and intracellular bacteria proliferation through its association with vacuolin A (VacA), a core component of host membrane microdomains at the MCV. Our data identify VacA as a central regulator of microdomain stability, independently of the other vacuolin isoforms, VacB and VacC. Collectively, our findings uncover a regulatory circuit in which DstA promotes *vacA* transcription and VacA accumulation at the MCV membrane, while VacA reciprocally restricts DstA nuclear translocation acting as a cytoplasmic anchor. This feedback loop couples transcriptional regulation with membrane composition homeostasis to fine-tune host anti-mycobacterial defence.

## Results

### DstA and DstB are host susceptibility factors, while DstC is a resistance factor

To determine the role of Dst proteins during the early phase of host defence, we first assessed the dynamics of transcription of the four Dst genes during infection of *Dd* with *Mm* WT using available RNA sequencing data ^43^. The genes encoding DstA, DstC and DstD are transiently upregulated at 1 hour post-infection (hpi), followed by a slight global downregulation or unchanged expression from 3 to 12 hpi compared to mock non-infected *Dd* (Supplementary Fig. 1a). To further determine their involvement during infection, we generated individual *dst* knockout (KO) cell lines using a single guide RNA approach to produce CRISPR/Cas9-mediated deletions (Fig. 1a-c and Supplementary Fig. 1b). We next established complemented cell lines stably expressing the corresponding GFP-tagged Dst proteins from a “safe haven” chromosomal locus (Fig. 1a-c). All cell lines were validated by DNA sequencing of the genomic region encompassing the target site and by western blot analysis (Fig. 1a-c). The *dstB*-KO cell line was validated exclusively by DNA sequencing due to the absence of specific antibody recognizing DstB (Fig. 1a-c and Supplementary Fig. 1b) ^44–47^. As shown in the Figure 1a-c, the KO cell lines lacked detectable DstA, DstC and DstD proteins compared to the WT strain, while the complemented cell lines expressed the corresponding GFP-Dst fusion proteins. We next characterized the impact of *dst* deletion on intracellular *Mm* growth. *Dd* cell lines were infected with bacterial luciferase-expressing *Mm*, and bacteria burden was monitored over time by measuring luminescence. Intracellular *Mm* growth in *dstD*-KO clones was comparable to that observed in WT cells (Fig. 1d). In contrast, bacteria growth was attenuated in *dstA-* and *dstB*-KO cells (Fig. 1e-f), whereas it was increased in *dstC*-KO cells (Fig. 1g). These phenotypes were confirmed using independent KO clones harbouring distinct deleterious mutations (Supplementary Fig. 1c-e). Re-expression of GFP-Dst in the corresponding KO cell lines rescued the observed phenotypes (Fig. 1e-g). Collectively, these data identify DstA and DstB as host susceptibility factors, while DstC functions as a resistance factor.

**Figure 1.**
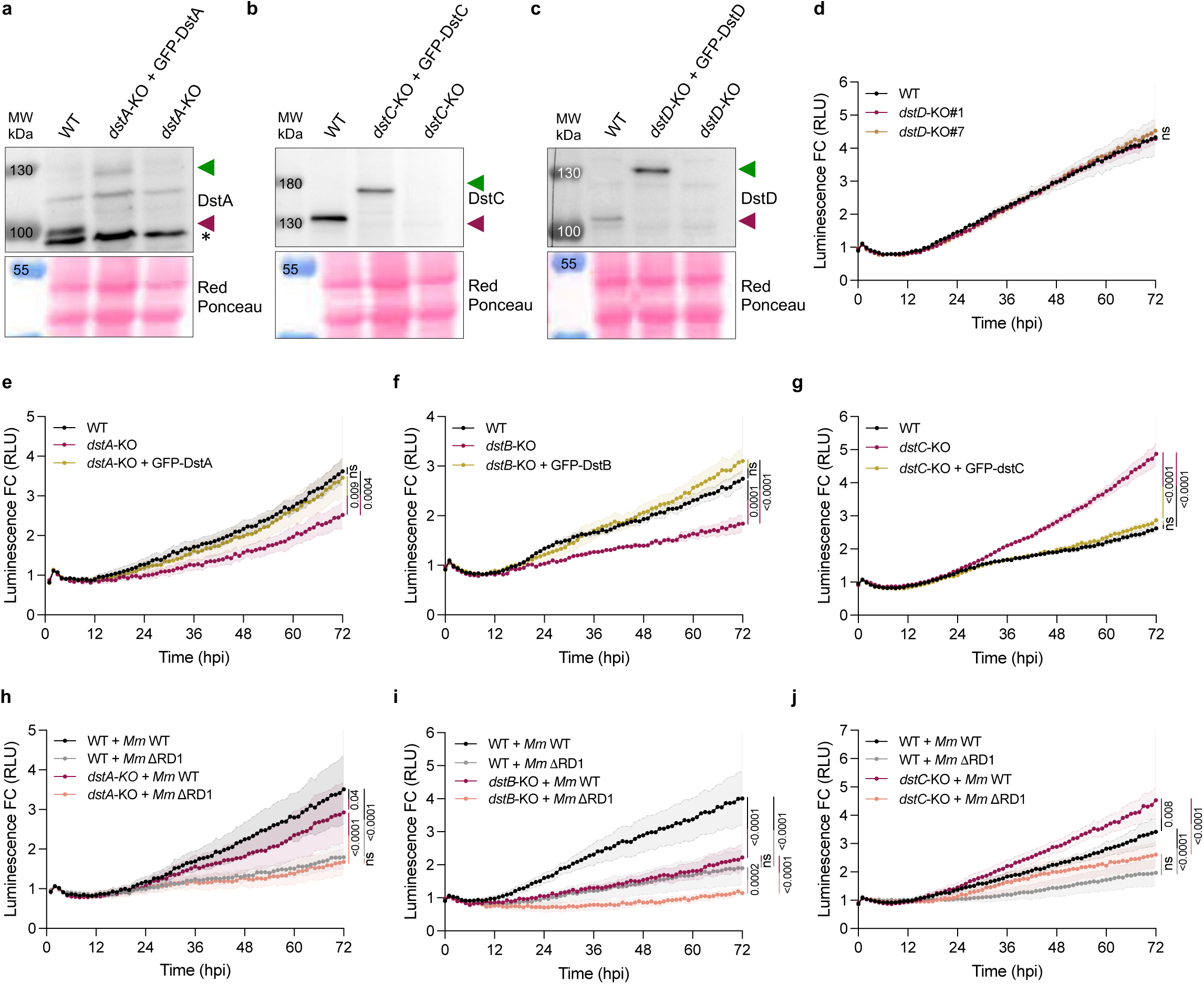
DstA, DstB and DstC impact *M. marinum* intracellular growth. **A-C)** Representative immunoblots of WT, *dst*-knockout (KO), and *dst*-KO expressing GFP-tagged-Dst proteins, using the indicated antibodies. The green arrow indicates the GFP-tagged protein, whilst the red arrow indicates the endogenous protein. * represents a non-specific band. **D)** *Dd* WT and two independent clones of *dstD*-KO cells were infected with bioluminescent *Mm* WT. Bioluminescence was measured for 72 hours (mean fold change ± SEM, *N* = 3, one-way ANOVA, Tukey’s multiple comparisons test). RLU, relative luminescence unit. **E-G)** *Dd* WT, *dst*-KO and *dst*-KO cells expressing GFP-Dst were infected with bioluminescent *Mm* WT. Bioluminescence was monitored over 72 hours (mean fold change ± SEM, *n* = 3, *N* ≥ 3, one-way ANOVA with Tukey’s multiple comparisons test). RLU, relative luminescence units. **H-J)** *Dd* WT and *dst*-KO cells were infected with bioluminescent *Mm* WT or ΔRD1. Bioluminescence was monitored over 72 hours (mean fold change ± SEM, *n* = 3, *N* = 3, one-way ANOVA, Tukey’s multiple comparisons test). RLU, relative luminescence units.

To investigate the influence of DstA, DstB and DstC on host responses to *Mm*–induced MCV damage, we next examined the intracellular growth of *Mm* ΔRD1, a strain lacking the Region of Difference 1 (RD1) which encodes the ESX-1 system required for membrane disruption and cytosolic access. Growth of this damage-deficient *Mm* strain in *dstA*-KO cells was comparable to that in WT *Dd* cells (Fig. 1h), indicating that the attenuation observed with *Mm* WT in *dstA*-KO cells depends on bacterial membrane-damaging activity. In contrast, the deletion of DstB and DstC resulted in attenuated and enhanced intracellular growth, respectively, even in the absence of ESX1-mediated damage (Fig. 1i-j). These findings suggest that the functions of DstB and DstC are not solely dependent on *Mm*-induced membrane damage but also involve additional susceptibility and resistance mechanisms.

### DstA, DstB and DstC control host responses during infection

To investigate host transcriptional programs regulated by DstA, DstB and DstC, we infected KO cell lines with GFP-expressing *Mm* and isolated infected cells (GFP-positive), and bystander cells (GFP-negative) by FACS, as previously described ^43^. RNAs were then extracted, and RNA sequencing was performed at 1, 6 and 24 hpi. Differentially expressed genes (DEGs) were identified by comparing infected *Dd dst*-KO cells to infected WT cells. This analysis revealed a marked increase in both downregulated and upregulated genes in KO cell lines during infection with a pronounced enrichment of downregulated genes at early timepoints (1 and 6 hpi) (Fig. 2a). These results indicate that DstA, DstB and DstC broadly promote transcriptional activation during infection, either directly or indirectly.

**Figure 2.**
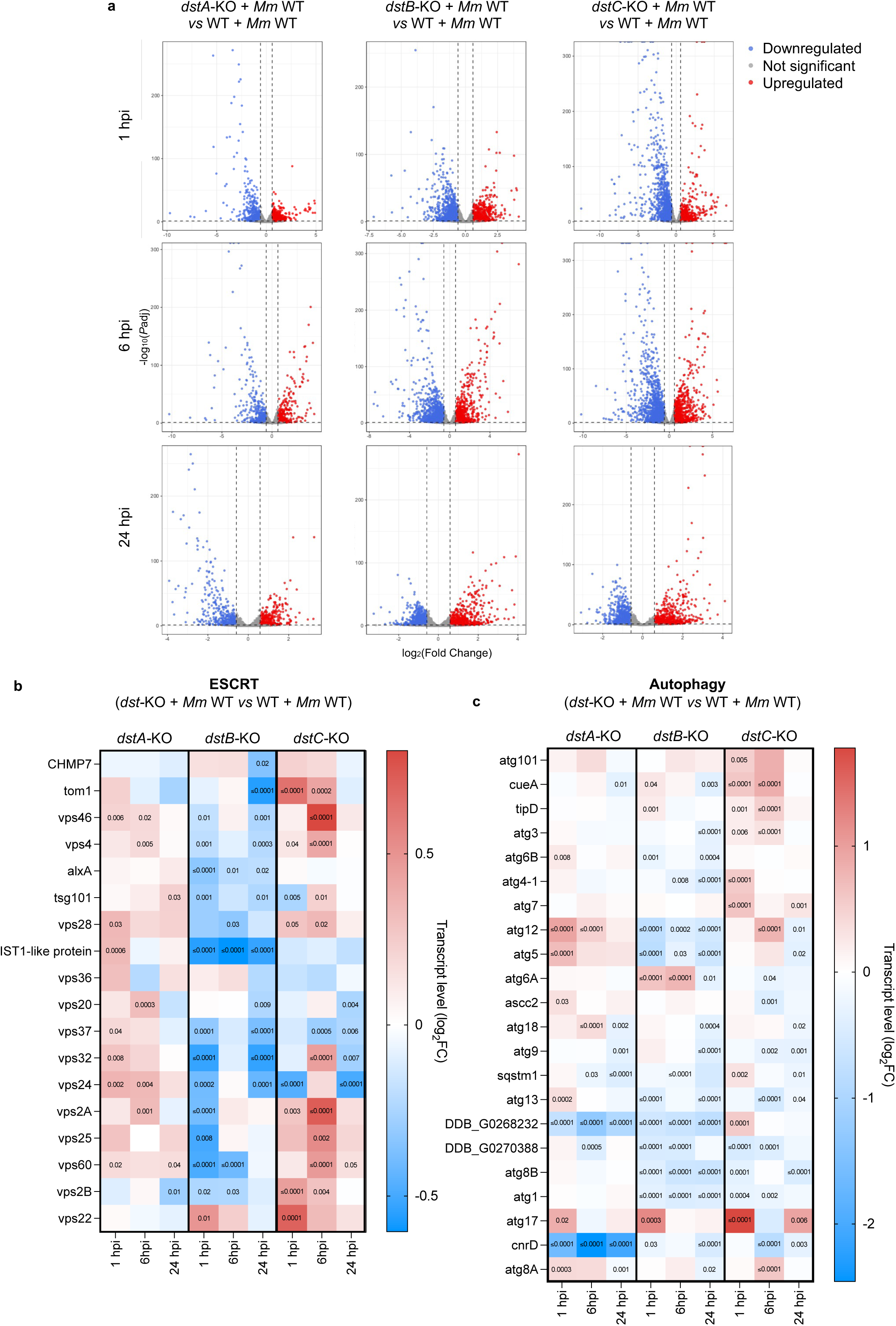
DstA, DstB and DstC control membrane repair responses during infection. **A)** Volcano plots of RNA-seq analysis. Selected threshold for statistical significance is *P*adj < 0.05.RNA-seq data of infected *dst*-KO cells are normalized to infected WT cells and is in log_2_ Fold Change (*N* = 3). Blue dots represent downregulated genes and red dots represent upregulated genes. **B-C)** Heatmaps showing RNA-seq transcriptome analysis for **B)** ESCRT- or **C)** autophagy-mediated repair genes expression. RNA-seq data from infected *dst*-KO cells are normalized to infected WT cells and is represented as log_2_ fold change (*N* = 3).

In *dstA*-KO cells, KEGG pathway enrichment analysis of the downregulated DEGs revealed persistent impact on genes related to lysosomal functions throughout the course of infection, including pathways associated with cysteine proteases, Rho GTPases, ATPases, peptidases and glucosaminidases (Supplementary Fig. 2). At later stages of infection, pathways related to fatty acid biosynthesis were also significantly enriched among the downregulated genes, including methylmalonyl-CoA mutase, acyl-CoA synthetase, acyl-CoA oxidases, malonyl-CoA decarboxylase (Supplementary Fig. 2). In contrast, the *dstB-* and *dstC*-KO cells exhibited a broader transcriptional alteration encompassing multiple functional categories. Notably, at 1 hpi, gene clusters associated with transcription and protein synthesis were strongly downregulated. In *dstB*-KO cells, additional downregulated pathways included central metabolism and cofactor biosynthesis, membrane dynamics and signalling pathways, lysosomal degradation, and host–pathogen interaction processes over the course of infection (Supplementary Fig. 2). Similarly, *dstC*-KO cells displayed downregulation of metabolic and biosynthetic processes, membrane dynamics and host defence pathways, as well as cytoskeleton-dependent transport mechanisms (Supplementary Fig. 2). Collectively, these results demonstrate that loss of DstA, DstB, or DstC leads to widespread downregulation of diverse host pathways during infection, supporting a central role for these factors in coordinating transcriptional responses to *Mm* invasion.

Once inside the MCV, *Mm* induces membrane damage, which is subsequently repaired through the coordinated action of ESCRT and Atg8-dependent autophagy machineries, thereby limiting bacteria escape to the cytosol. We therefore investigated whether Dst transcription factors modulate the engagement of these pathways during infection. Focusing on ESCRT-related genes, we observed a marked downregulation in *dstB*-KO cells at 1, 6 and 24 hpi compared to infected WT cells (Fig. 2b). In contrast, these genes were predominantly upregulated in *dstA-* and *dstC*-KO cells at 1 and 6 hpi (Fig. 2b). To determine whether these transcriptional signatures are infection-specific, we performed RNA sequencing on *dstA-*, *dstB-* and *dstC*-KO cells under steady state conditions (Supplementary Fig. 3). Notably, ESCRT-related genes were only modestly differentially expressed in *dstA-* and *dstB*-KO cells at steady state compared to WT cells (Supplementary Fig. 3a), indicating that their regulation is largely infection-dependent. In contrast, these genes were already upregulated at steady state in *dstC*-KO cells (Supplementary Fig. 3a), showing that the ESCRT pathway is intrinsically modulated in this background. Similarly, autophagy-related genes were slightly downregulated in infected *dstB*-KO cells compared to infected WT (Fig. 2c), whereas no significant changes were observed at steady state (Supplementary Fig. 3b), further supporting an infection-specific regulation. Together, these findings suggest that DstA, DstB, and DstC exert distinct and infection-dependent control over ESCRT- and autophagy-mediated membrane repair, shaping the host response to *Mm*-induced phagosomal damage.

### DstA, DstB and DstC regulate host vacuolar responses during infection

We next sought to determine whether *Mm* escapes prematurely into the cytosol or instead remains confined within the MCV as a consequence of metabolic limitation, or restriction by antibacterial mechanisms operating either within the vacuole or in the cytosol. To address this, we monitored the association of the lipid droplet protein perilipin (Plin) with *Mm* upon cytosolic exposure, as Plin recruitment is commonly used as a reporter of bacterial escape from the MCV ^48^. In WT cells, association of mCherry-Plin with *Mm* increased from 6 hpi onwards, confirming the progressive escape of *Mm* to the cytosol (Fig. 3a-b). A similar pattern was observed in *dstB*-KO cells (Fig. 3a-b), with no detectable defect in Plin coating, as mCherry-Plin covered a comparable area of the bacterial surface in these cells (Fig. 3d). In contrast, mCherry-Plin recruitment to *Mm* was strongly reduced in *dstA-* and *dstC*-KO cells (Fig. 3a-b), indicating that *Mm* remains confined within the MCV in these genetic backgrounds. Consistent with these observations, the attenuated bacteria growth and low level of cytosolic bacteria in *dstA*-KO cells indicate that *Mm* remains confined in the MCV and thereby attenuated. In contrast, the enhanced bacterial growth and low level of cytosolic bacteria observed in *dstC*-KO cells suggest that *Mm* remains confined within the MCV but exhibits an improved intravacuolar fitness. Finally, in *dstB*-KO cells, the attenuated intracellular growth combined with unaffected cytosolic escape suggest a stronger restriction of *Mm* in the cytosol. During infection with the *Mm* ΔRD1 mutant strain, mCherry-Plin recruitment in all KO cell lines was comparable to that observed in WT cells (Fig. 3c), showing that in absence of ESX-1-mediated damage, MCV confinement is maintained. In line with previous results, this suggests that none of the *dst*-KO cells promote premature *Mm* cytosolic escape.

**Figure 3.**
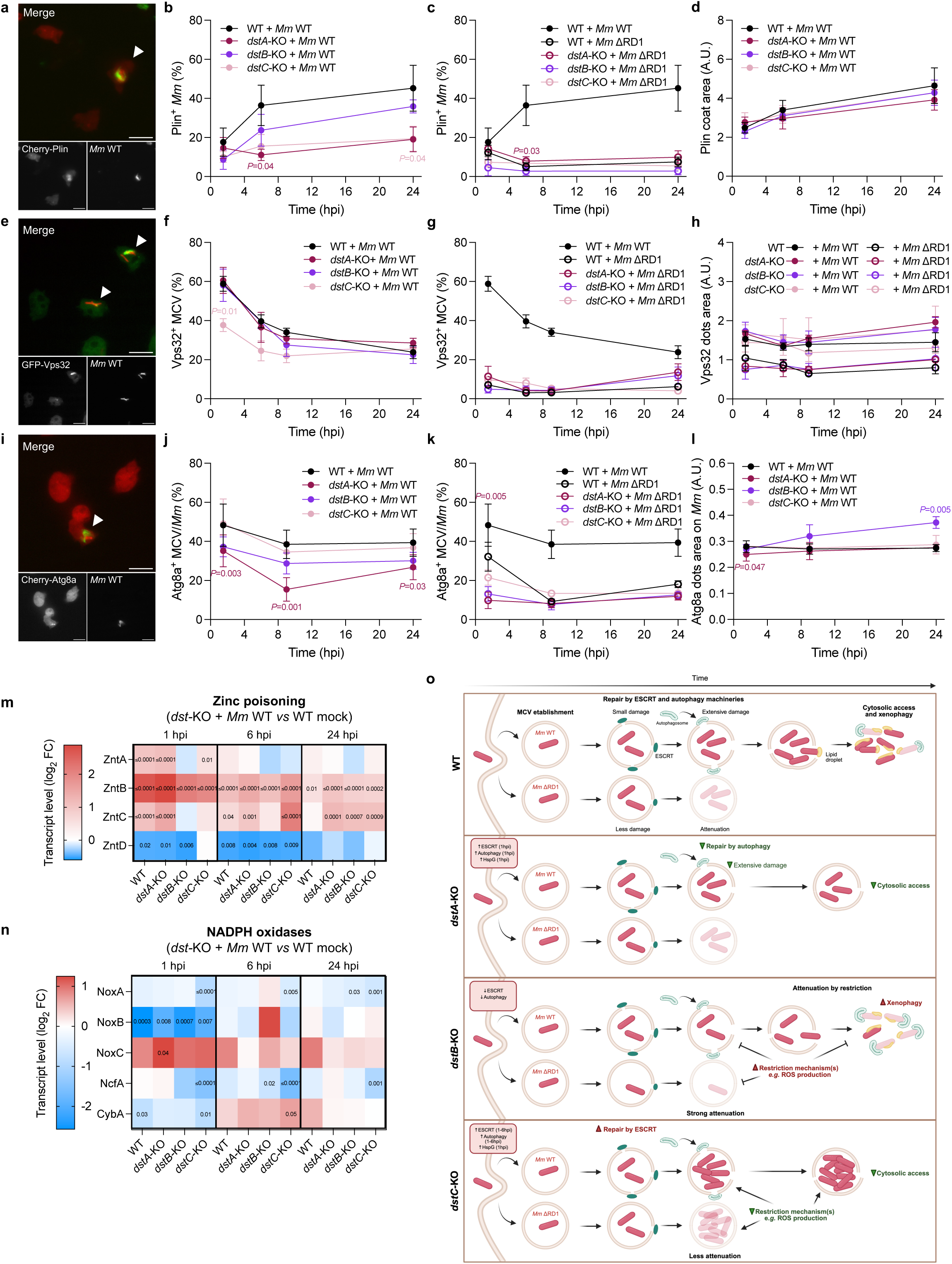
DstA, DstB and DstC regulate bacterial escape by modulating ESCRT and autophagy pathways. **A-D)** *Dd* WT and *dst*-KO cells expressing mCherry-Plin were infected with GFP-expressing *Mm* WT or ΔRD1 mutant and imaged live by high-content microscopy. **A)** Representative images at 9 hpi. White arrow indicates infected cells. Scale bars: 10 μm. **B-C)** Quantification of A, showing the percentage of infected cells with Plin-*Mm* colocalization or **D)** the area of Plin labelling on *Mm* (mean ± SD, *n* = 3; *N* = 4, two-way ANOVA with Tukey’s multiple comparisons test). **E-H)** *Dd* WT and *dst*-KO cells expressing GFP-Vps32 were infected with mCherry-expressing *Mm* WT or ΔRD1 mutant and imaged live by high-content microscopy. **E)** Representative images at 1 hpi. White arrow indicates infected cells. Scale bars: 10 μm. **F-G)** Quantification of E, showing the percentage of infected cells with Vps32-*Mm* colocalization or **H)** the area of Vps32 labelling on *Mm* (mean ± SD, *n* = 3; *N* ≥ 3, two-way ANOVA with Tukey’s multiple comparisons test). **I-L)** *Dd* WT and *dst*-KO cells expressing mCherry-Atg8a were infected with GFP-expressing *Mm* WT or ΔRD1 mutant and imaged live by high-content microscopy. **I)** Representative images at 1 hpi. White arrow indicates infected cells. Scale bars: 10 μm. **J-K)** Quantification of I, showing the percentage of MCV with Atg8a labelling or **L)** the area of Atg8a labelling on *Mm* (mean ± SD, *n* = 3; *N* ≥ 3, two-way ANOVA with Tukey’s multiple comparisons test). **M-N)** Heatmaps showing RNA-seq transcriptome analysis for **M)** Zinc poisoning or **N)** NADPH oxidases-related genes expression. RNA-seq data from infected *dst*-KO cells are normalized to non-infected WT cells and is represented as log_2_ Fold Change. **O)** Schematic model depicting the fate of infection in the different *dst*-KO cells. MCV, *Mycobacterium*-containing vacuole; ROS, reactive oxygen species.

To determine whether membrane repair mechanisms are involved in the observed phenotypes, we quantified the recruitment of the ESCRT-III component GFP-Vps32 and the autophagy/Atg8ylation reporter mCherry-Atg8a to the MCV. Early recruitment of GFP-Vps32 to the MCV in WT cells indicates activation of ESCRT-mediated membrane repair in response to EsxA-induced small damage (Fig. 3e-f) ^36,40^. ESCRT engagement was impaired in *dstC*-KO cells at early time points, as indicated by a reduced GFP-Vps32 localization to the MCV compared to WT cells (Fig. 3f). In contrast, recruitment of GFP-Vps32 to ΔRD1-containing vacuoles was not affected (Fig. 3g). However, GFP-Vps32 structures associated with the damage-deficient mutant were enlarged, reaching size comparable to those observed during infection with *Mm* WT (Fig. 3h). Given that *Mm* remains confined in the MCV in *dstC*-KO cells, these results suggest a more efficient ESCRT-mediated membrane repair response.

While the ESCRT machinery promotes repair of small membrane lesions, extensive damage leads to the recruitment of the Atg8-dependent autophagy machinery (Fig. 3i-j) ^37^. The recruitment of mCherry-Atg8a to the MCV was strongly reduced in *dstA*-KO cells over the infection with *Mm* WT, and at 1 hpi during infection with ΔRD1 mutant (Fig. 3j-k). Furthermore, mCherry-Atg8a dots at the MCV in *dstA*-KO cells were smaller than in WT cells at 1 hpi (Fig. 3l). Together with the observed confinement of *Mm* within the MCV in *dstA*-KO cells and the unaffected ESCRT engagement, these data suggest two possibilities: either damage to the MCV membrane is efficiently repaired by the ESCRT machinery, thereby reducing the requirement for Atg8-dependent autophagic repair, or *Mm* induces less membrane damage, potantially due to altered vacuolar membrane composition that renders the MCV more resistance to EsxA-mediated membranolysis, consequently limiting the recruitment of autophagy machinery. In *dstB*-KO cells, mCherry-Atg8a recruitment to the MCV was comparable to that in WT cells, however the mCherry-Atg8a dots were larger at 24 hpi (Fig. 3g-h, j). Given that the absence of DstB leads to cytosolic escape and restriction of *Mm*, these data suggest that the autophagy-mediated repair is not impaired, but rather that xenophagy or other antimicrobial defence is enhanced.

We hypothesized that xenophagy might contribute to the cytosolic restriction of *Mm* in *dstB*-KO cells. Given that xenophagy is triggered downstream of the disruption of the MCV membrane, leading to Ca^2+^ leakage and bacteria exposure to the cytosol ^49^, the observed attenuated growth of *Mm* ΔRD1 in this cell line compared to WT *Dd* indicates that additional damage-independent antibacterial mechanisms are involved.

An opposite observation was made in *dstC*-KO cells, where *Mm* ΔRD1 growth was enhanced, showing that damage-independent antibacterial mechanisms are impaired in this genetic background. To further investigate this, we analysed the transcriptional expression of genes involved in other antimicrobial pathways, including the generation of ROS and metal poisoning ^24,28,50,51^. To determine whether a specific transcriptional signature is driven both by the infection and the genetic background, we assessed the enrichment of these defence-related genes during infection compared to uninfected WT *Dd*. Pathway analysis revealed that transcription of genes involved in neither zinc nor copper poisoning pathways were significantly altered in these mutants (Fig. 3m and Supplementary Fig. 4a). On the other hand, interestingly, NADPH oxidases were upregulated in *dstB*-KO cells and downregulated in *dstC*-KO cells at 6 hpi (Fig. 3n), consistent with ROS-dependent antibacterial process being reinforced in *dstB*-KO cells and diminished in *dstC*-KO cells. Intriguingly, members of the HspG family, which belong to small Heat Shock Proteins (sHSPs), were strongly downregulated at the early stages of infection in WT cells (Supplementary Fig. 4b). In mammals, sHSPs are known for their anti-oxidant and anti-apoptotic functions, as well as for maintaining cytoskeletal integrity ^52–55^. In marked contrast to the infection in WT cells, these genes were only weakly downregulated during infection in *dstA-* and *dstC*-KO cells at the first timepoint, leading to the hypothesis that it might be implicated in counteracting oxidative stress or preventing cell death. Altogether, these results indicate that DstA, DstB and DstC sequentially regulate distinct restriction pathways during infection (Fig. 3o).

### *Mm*-induced damage drives early nuclear translocation of DstA followed by cytosolic relocalization of DstB and DstC

To assess the dynamic nucleo-cytoplasmic redistribution of DstA, DstB and DstC during infection, *dst*-KO cells expressing the corresponding GFP-Dst constructs and mCherry-Histone 2B (H2B) were used to quantify nuclear *versus* cytosolic localization (Supplementary Fig. 5a). Under steady state conditions, GFP-DstA localized to both the cytosol and nucleus, whereas GFP-DstB and GFP-DstC were predominantly nuclear (Supplementary Fig. 5b). Following infection with blue fluorescent *Mm* pTEC18, GFP-DstA translocated to and accumulated inside the nucleus of infected cells, while remaining cytosolic in uninfected and bystander cells (Fig. 4a). This nuclear localization progressively decreased over the course of infection (Fig. 4a-c) and was not associated with a decrease in DstA protein levels (Supplementary Fig. 5c-d), indicating a transient nuclear translocation of DstA at very early stages of infection. Importantly, infection with *Mm* ΔRD1 mutant did not impact GFP-DstA localization, which was similar to that observed in non-infected and bystander cells (Fig. 4a-c). To determine the precise timing of this nuclear translocation, *dstA*-KO cells expressing GFP-DstA were incubated with *Mm* WT or ΔRD1 pTEC18 in order to capture early events shortly after phagocytosis. GFP-DstA enrichment in the nucleus was detected at the earliest stages of infection with *Mm* WT (Fig. 4d), but not with ΔRD1 mutant (Supplementary Fig. 5e), attesting of a rapid and damage-dependent nuclear translocation. Of note, a small fraction of GFP-DstA forms puncta in close proximity to *Mm* WT after phagocytosis (Fig. 4d). Following infection with *Mm* WT, nuclear levels of GFP-DstB gradually decreased, demonstrating progressive relocalisation of GFP-DstB to the cytosol from 4 hpi onward (Fig. 4a, e-f). Interestingly, this phenomenon was also observed in bystander cells, whereas GFP-DstB remained predominantly nuclear during infection with *Mm* ΔRD1 mutant (Fig. 4a, e-f). A similar pattern was found with GFP-DstC (Fig. 4a, g-h), suggesting a shared mechanism involving damage-dependent relocalization to the cytosol of infected cells that is also communicated to neighbouring non-infected cells. Moreover, the phosphorylation level of DstC increased during infection with *Mm* ΔRD1 mutant, whereas it remained unchanged during infection with *Mm* WT (Supplementary Fig. 5f-g). This suggests that MCV membrane damage decreases DstC phosphorylation, which may be linked to the relocalization to the cytosol of the dephosphorylated form. Altogether, these results reveal a dynamic redistribution of the Dst transcription factors, in which *Mm*-induced damage drives early nuclear translocation of DstA, followed by later cytosolic relocalization of DstB and DstC (Fig. 4i). The response of DstA is restricted to infected cells, whereas the signalling triggering DstB and DstC relocalization propagates beyond infected cells (Fig. 4i).

**Figure 4.**
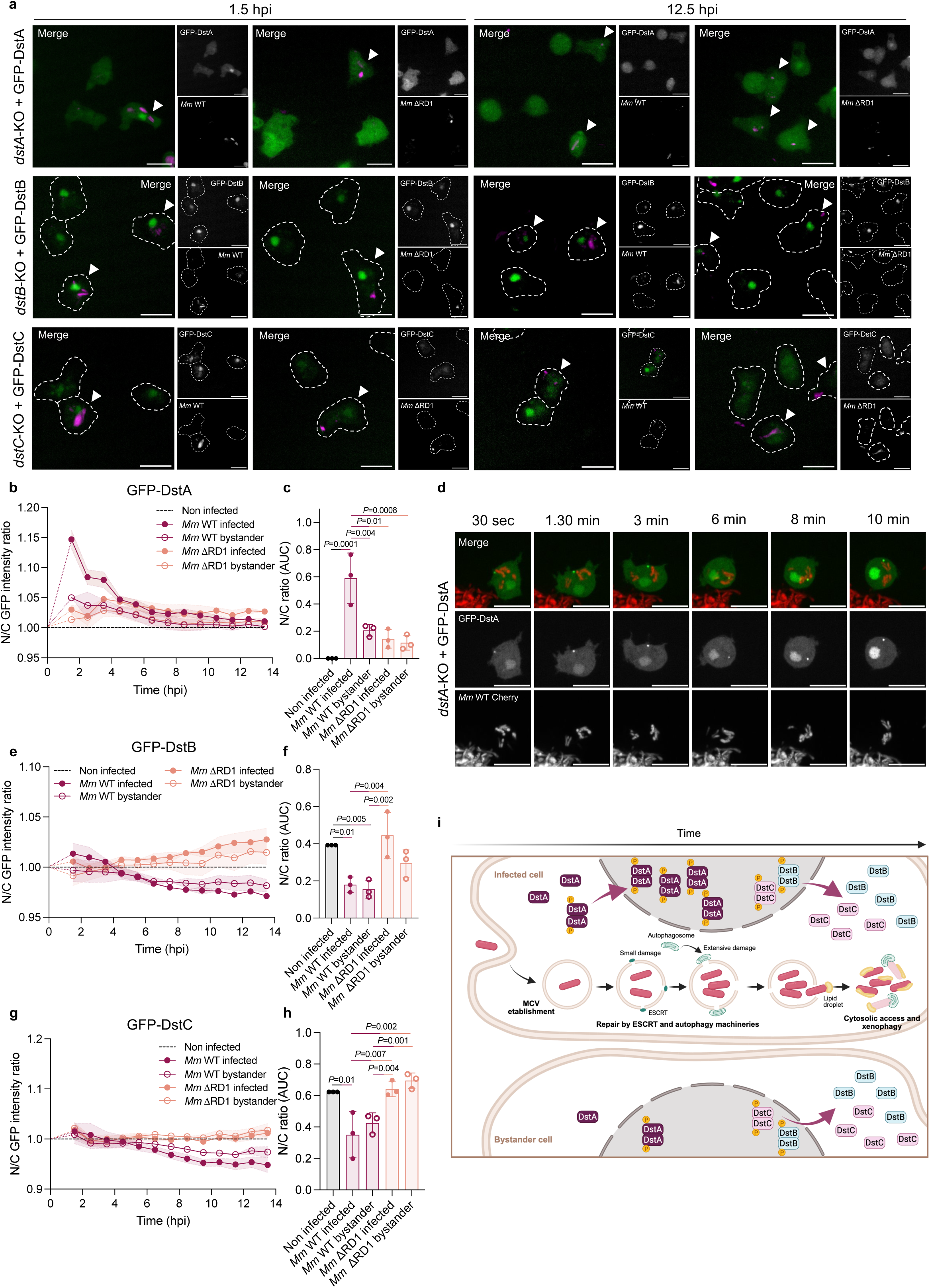
*Mm*-induced damage drives early nuclear translocation of DstA followed by cytosolic relocalization of DstB and DstC. **A-C)** *Dd dst*-KO cells expressing GFP-Dst and mCherry-H2B were infected with blue-fluorescent pTEC18 *Mm* WT or ΔRD1 mutant and imaged live by high-content microscopy. **A)** Representative live images at 1.5 and 12 hpi. White arrow indicates infected cells. Scale bars: 10 μm. **B)** Quantification of A, showing the nuclear-to-cytosolic GFP-DstA intensity ratio in infected and bystander cells, normalized to the ratio in non-infected cells (mean ± SEM, *n* = 3; *N* = 3). N/C; nucleo-cytoplasmic ratio. **C)** Aera under the curve of B (mean ± SD, *N* = 3, One-way ANOVA). **D)** Live microscopy of *Dd dstA*-KO cells with mCherry-expressing *Mm* WT shortly after phagocytosis. Following incubation with *Mm,* infected cells was imaged for 10 min. Scale bars: 10 μm. **E)** Quantification of A, showing the nuclear-to-cytosolic GFP-DstB intensity ratio in infected and bystander cells, normalized to the ratio in non-infected cells (mean ± SEM, *n* = 3; *N* = 3). **F)** Aera under the curve of E (mean ± SD, *N* = 3, One-way ANOVA). **G)** Quantification of A, showing the nuclear-to-cytosolic GFP-DstC intensity ratio in infected and bystander cells, normalized to the ratio in non-infected cells (mean ± SEM, *n* = 3; *N* = 3). **H)** Aera under the curve of G (mean ± SD, *N* = 3, One-way ANOVA). **I)** Schematic model depicting the translocation of Dst proteins during infection.

### *Mm*-induced damage is required for DstA, DstB and DstC translocation

L-leucyl-L-leucine methyl ester (LLOMe) is an endolysosomal membrane-disrupting agent that induces reversible, synchronous and homogenous damage, phenocopying several features of EsxA-induced membrane damage ^42^. To validate the damage-dependent nature of DstA translocation, GFP-DstA nuclear localization was quantified upon LLOMe treatment in *dstA*-KO cells expressing GFP-DstA. LLOMe treatment triggered rapid nuclear translocation compared to untreated cells, peaking at 20 minutes post-treatment (Fig. 5a-c). Importantly, as already shown, GFP-Vps32 recruitment to sites of membrane damage was rapid and transient (Supplementary Fig. 6a-b) ^42^, whereas nuclear translocation of GFP-DstA was more sustained over time (Fig. 5a-c). These results confirm that membrane damage is sufficient to trigger DstA nuclear translocation. In addition, this response was abolished when cells were pre-treated with methyl-β-cyclodextrin (MβCD, Fig. 5d-f and Supplementary Fig. 6c-d), which extracts sterols from membranes and disrupts cholesterol-rich lipid raft domains (microdomains), thereby altering membrane susceptibility to LLOMe- and EsxA-triggered damage ^42,56^. Indeed, although mechanistically distinct, the activity of both membrane disrupting agents requires membrane microdomains ^42^. A similar effect was observed during infection in presence of MβCD, where DstA nuclear translocation was abolished, phenocopying infection with the *Mm* ΔRD1 mutant (Fig. 5g-i). Cytosolic relocalization of DstB and DstC was likewise reduced, although not completely abolished, in the presence of MβCD compared to non-infected cells (Fig. 5g, j-m). Overall, during infection, early DstA nuclear translocation strictly requires an organized MCV membrane and *Mm*-induced damage, whereas DstB and DstC localization is influenced by both damage-dependent and damage-independent signals, highlighting broader regulatory roles of these Dst transcription factors.

**Figure 5.**
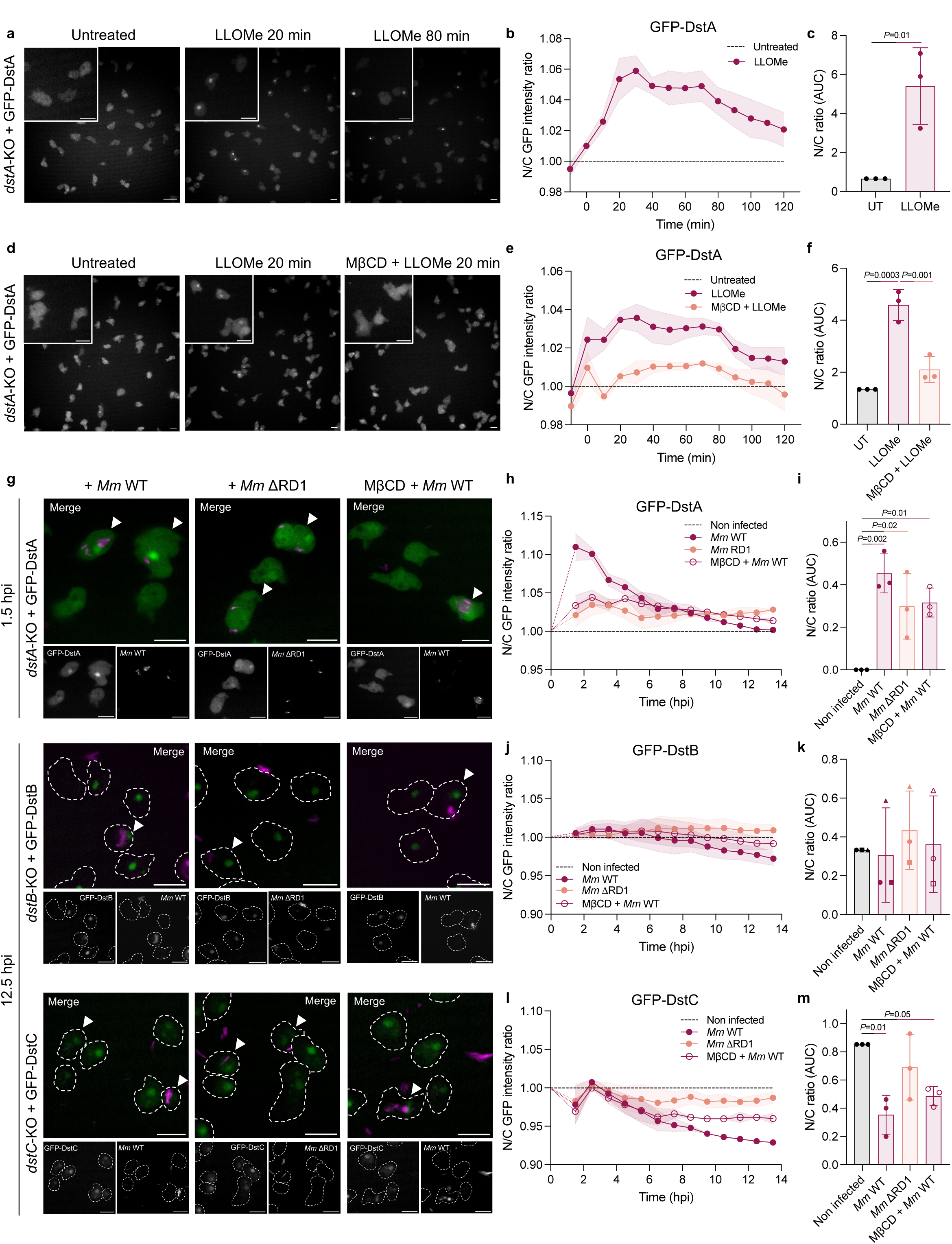
*Mm*-induced damage is required for DstA, DstB and DstC translocation. **A-C)** *Dd dstA*-KO cells expressing GFP-DstA and mCherry-H2B were treated with 4.5 mM of LLOMe or left untreated. **A)** Representative live images showing GFP-DstA localization before treatment and 20 min after LLOMe addition. Scale bars: 10 μm. **B)** Quantification of A, showing the nuclear-to-cytosolic GFP-DstA intensity ratio, normalized to the ratio in untreated cells (mean ± SEM, *n* = 2; *N* = 3). **C)** Aera under the curve of B (mean ± SD, *N* = 3, One-way ANOVA). N/C; nucleo-cytoplasmic ratio. **D-F)** *Dd dstA*-KO cells expressing GFP-DstA and mCherry-H2B pretreated or not with 2 mM of MβCD, were then treated with 4.5 mM LLOMe. **D)** Representative live images showing GFP-DstA localization before treatment and 20 min after LLOMe addition. Scale bars: 10 μm. **E)** Quantification of D, showing the nuclear-to-cytosolic GFP-DstA intensity ratio, normalized to the ratio in untreated cells (mean ± SEM, *n* = 2; *N* = 3). **F)** Aera under the curve of E (mean ± SD, *N* = 3, One-way ANOVA). **G-M)** *Dd dst*-KO cells expressing GFP-Dst construct and mCherry-H2B pretreated or not with 2 mM of MβCD, were infected with blue-fluorescent pTEC18 *Mm* WT or ΔRD1 mutant and imaged live by high-content microscopy. **G)** Representative live images. White arrow indicates infected cells. Scale bars: 10 μm. **H)** Quantification of G, showing the nuclear-to-cytosolic GFP-DstA intensity ratio in infected and bystander cells, normalized to the ratio in non-infected cells (mean ± SEM, *n* = 3; *N* = 3). **I)** Aera under the curve of H (mean ± SD, *N* = 3, One-way ANOVA). **J)** Quantification of G, showing the nuclear-to-cytosolic GFP-DstB intensity ratio in infected and bystander cells, normalized to the ratio in non-infected cells (mean ± SEM, *n* = 3; *N* = 3). **K)** Aera under the curve of J (mean ± SD, *N* = 3, One-way ANOVA). **L)** Quantification of G, showing the nuclear-to-cytosolic GFP-DstC intensity ratio in infected and bystander cells, normalized to the ratio in non-infected cells (mean ± SEM, *n* = 3; *N* = 3). **M)** Aera under the curve of L (mean ± SD, *N* = 3, One-way ANOVA).

### DstA, DstB and DstC physically associate with signalling proteins

To determine the interaction partners of these transcription factors, protein lysates from infected *dstA-*, *dstB-* and *dstC*-KO cells expressing the corresponding GFP-tagged Dst proteins were subjected to magnetic GFP-Trap purification at 1 hpi, followed by mass spectrometry analysis. Of note, this proteomic analysis identified phosphorylation of DstA at Tyr948, whereas no phosphorylation was detected for DstB and DstC, consistent with the previously observed translocation patterns at 1 hpi. Importantly, GFP-DstA, GFP-DstB and GFP-DstC were found to associate with multiple Gα subunits of the heterotrimeric G protein signalling pathways. In particular, GpaA (Gα1), GpaD (Gα4), GpaE (Gα5) and GpaL (Gα12) were identified as shared interactors of all three Dst proteins, while GpaI (Gα9) was specifically associated with DstA (Fig. 6a-b). To gain further insight into the role of G protein signalling during infection, we examined the transcription of genes encoding Gα subunits, as well as the Gβ (gpbA) and the Gγ (gpgA) subunits (Fig6C). Surprisingly, the expression of most Gα subunits remained unchanged at 1 hpi compared to non-infected WT *Dd*, but was upregulated between 3 to 12 hpi, followed by a downregulation from 24 to 48 hpi. This profile was not observed for *gpaD*, which displayed an opposite trend, with stable expression at early time points and an increased expression at later stages. Intriguingly, this subunit is a common interactor of DstA, DstB and DstC and was found to be upregulated during infection with the *Mm* ΔRD1 mutant compared to *Mm* WT (Fig. 6b, d). However, no specific transcriptional regulation of these Gα subunit was observed in the infected *dst*-KO cells (Supplementary Fig. 7a). Although the precise role of Gα protein signalling in the context of mycobacterial infection remains to be elucidated, this distinct expression signature and physical interaction with DstA, DstB and DstC highlights a potentially important contribution during infection.

**Figure 6.**
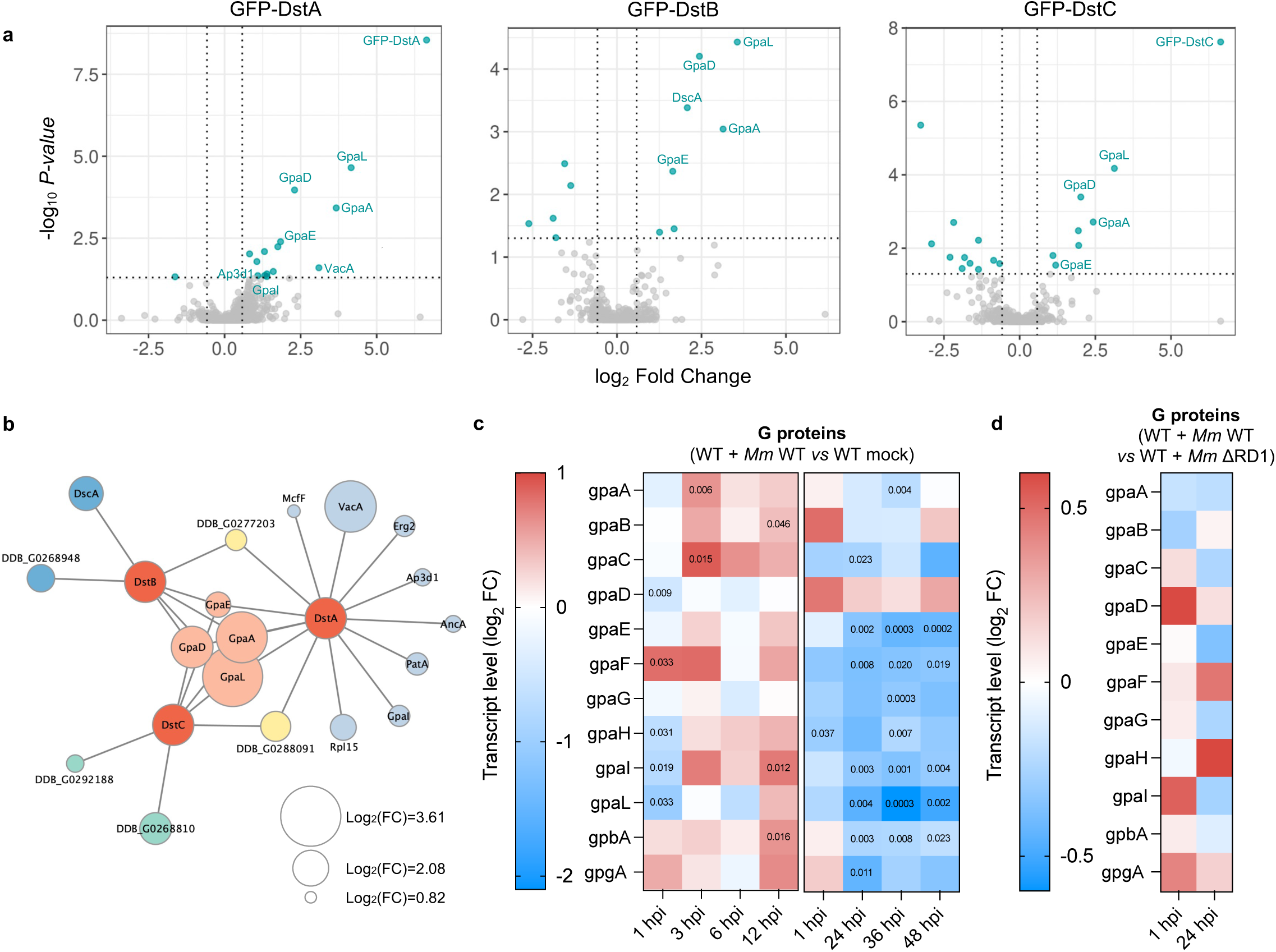
DstA, DstB and DstC interact with specific Gα subunits during infection. **A)** Pie charts representing the number of proteins detected in GFP-DstA, GFP-DstB and GFP-DstC pulldown, classified as significantly enriched or depleted (blue), or not significantly changed (grey) relative to GFP control. Selected threshold for statistical significance is log2 Ratio ≥ 0.58 and p-value ≤ 0.05. **B)** Protein-protein interaction network of DstA-, DstB-, DstC-associated proteins at 1 hpi according to A. Node size is proportional to log_2_ fold change relative to the GFP control condition. Node sizes for DstA, DstB and DstC were not adjusted. **C-D)** Heatmaps showing RNA-seq transcriptome analysis for *Gα* (*gpa*), *Gβ* (*gpbA*) and *Gγ* (*gpgA*) subunits expression. **C)** RNA-seq data from *Dd* WT infected with *Mm* WT are normalized to non-infected WT cells, and is represented as log_2_ fold change. **D)** RNA-seq data from *Dd* WT infected with *Mm* ΔRD1 are normalized to *Dd* WT infected with *Mm* WT, and is represented as log_2_ fold change.

In addition to shared interactors, each Dst protein displayed specific protein partners. DstA interacts with two mitochondrial substrate carrier family protein McfF and AncA (Fig. 6b). DstA also associates with the Adaptor-related Protein complex 3, Delta subunit 1 (Ap3d1), a component of the AP-3 adaptor complex involved in trafficking of cargo proteins from the trans-Golgi network and/or endosomes to lysosomes or lysosome-related organelles ^57^, as well as PatA, a Ca^2+^-induced plasma membrane ATPase ^58^. Interestingly, DstA interacts with vacuolin A (VacA), a functional flotillin homolog and evolutionarily conserved microdomain stabilizer at the endolysosomal and MCV membrane ^42,59^, and with Erg2, an isomerase involved in ergosterol biosynthesis. Ergosterol plays a role analogous to mammalian cholesterol and contributes to the organization of membrane microdomains. These specific interactions suggest potential functional specialization related to lysosomal trafficking and membrane microdomain organization. In addition, one specific partner of DstB is DscA, a cytosolic H-type lectin, functionally homologous to galectins, and participating in recognition of cytosolically exposed mycobacterial glycolipids ^60^. This interaction may contribute to DstB-dependent intracellular signalling after detection of cytosolic *Mm*. Interestingly, DscA is the only isoform of the four-member discoidin family that is downregulated in *dstB*-KO cells at 1 hpi and more markedly at 6 hpi (Supplementary Fig. 7b), a timepoint at which *Mm* begins to escape to the cytosol (Fig. 3b). This interaction analysis identifies functional partners directly involved in host cell-autonomous defence pathways.

### DstA selectively regulates VacA expression and its accumulation at the MCV

VacA, VacB, and VacC form hetero-oligomers and behave as integral membrane proteins that collectively coat the MCV membrane from 1 hpi onward ^42^. At the early stages of infection, DstA specifically interacts with VacA, whereas no interaction was detected with VacB and VacC. Given that no vacuolin isoform-specific association with the MCV was observed during infection ^42^, we investigated whether DstA transcriptionally regulates vacuolin expression during infection. In infected *dstA*-KO cells, *vacA* mRNA expression was downregulated at 1, 6 and 24 hpi, whereas *vacB* and *vacC* expression remained unchanged compared to infected WT cells (Fig. 7a-c). As *vacA* expression was also reduced at steady state (Supplementary Fig. 8a), we assessed the impact of infection on its expression in this background by comparing mRNA levels between infected and mock non-infected *dst*-KO cells. Interestingly, *vacA* was strongly downregulated at 1 hpi, and then its expression was progressively recovered over time, similar to the pattern of expression observed in infected WT cells (Supplementary Fig. 8b). Together, these results indicate that DstA selectively promotes *vacA* expression, while infection retains the ability to induce *vacA* upregulation even in DstA-deficient cells. VacA protein levels at early stages of infection were assessed in FACS-sorted cells infected with mCherry-expressing *Mm*. As previously described, VacA protein levels gradually accumulated over the course of infection in WT cells (Fig. 7d-e) ^42^. Corroborating the mRNA expression data, VacA protein expression was diminished in *dstA*-KO cells, with a more pronounced effect at 9 hpi (Fig. 7d-e), indicating that DstA positively regulates VacA expression. These results suggest that DstA contributes to the transcriptional control of vacuolar damage response pathways through selective regulation of the microdomain stabilizer VacA.

**Figure 7.**
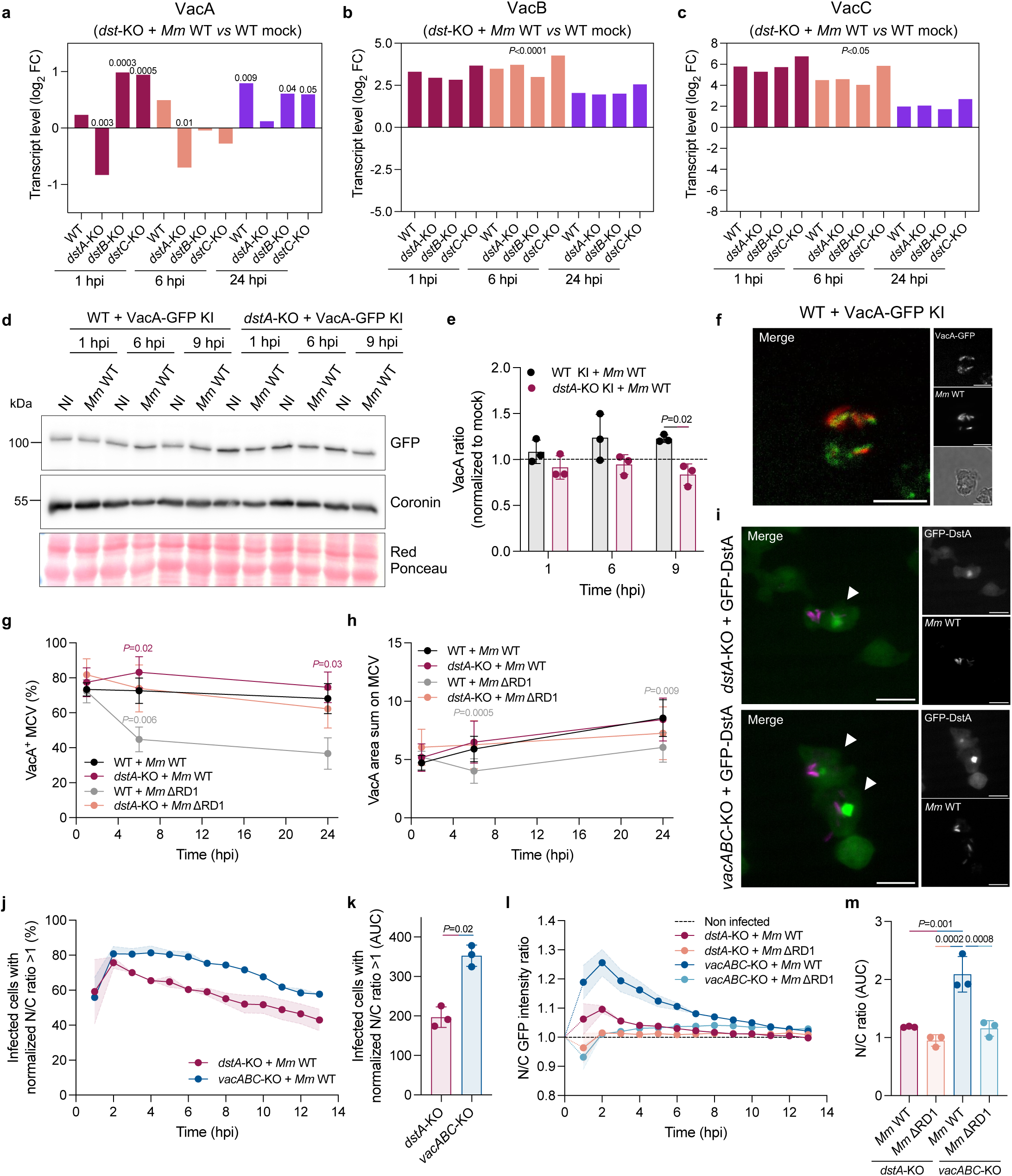
DstA and VacA form a regulatory circuit coordinating MCV integrity and transcriptional activity. **A-C)** RNA-seq analysis of *vacuolin* expression in FACS-sorted *Dd* WT or *dst*-KO cells infected with GFP-expressing *Mm* at the indicated hours post-infection (hpi). Fold changes in **A)** *vacA*, **B)** *vacB* and **C)** *vacC* transcript levels are shown relative to non-infected WT cells (mock, *N* = 3). **D)** Representative immunoblot of VacA-GFP knock-in (KI) in WT and *dstA*-KO cells infected with WT *Mm* or under non-infected mock conditions, using the indicated antibodies. Infected cells were sorted by FACS prior to analysis. **E)** Quantification of VacA-GFP represented in D) normalized to Coronin and mock condition (dashed line, mean ± SD, *N* = 3, two-way ANOVA with Tukey’s multiple comparisons test). **F-H)** VacA-GFP KI in *Dd* WT and *dstA*-KO cells were infected with mCherry-expressing *Mm* WT or ΔRD1 mutant and imaged live by high-content microscopy. **F)** Representative images of infected *Dd* WT VacA-GFP KI at 1 hpi. Scale bars: 10 μm. **G-H)** Quantification of F, showing the percentage of infected cells with VacA-*Mm* colocalization or **H)** the area of VacA labelling on *Mm* (mean ± SD, *n* = 3; *N* = 4, two-way ANOVA with Tukey’s multiple comparisons test). **I-M)** *Dd dstA*-KO and *vacABC*-KO cells expressing GFP-DstA and mCherry-H2B were infected with blue-fluorescent pTEC18 *Mm* WT or ΔRD1 mutant and imaged live by high-content microscopy. **I)** Representative live images at 1 hpi. White arrow indicates infected cells. Scale bars: 10 μm. **J)** Quantification of I, showing the percentage of infected cells exhibiting a nuclear-to-cytosolic GFP-DstA intensity ratio > 1, normalized to the corresponding ratio in non-infected cells (mean ± SEM, *n* = 3; *N* = 3). N/C; nucleo-cytoplasmic ratio. **K)** Aera under the curve of J (mean ± SD, *N* = 3, One-way ANOVA). **L)** Quantification of I, showing the nuclear-to-cytosolic GFP-DstA intensity ratio in infected cells, normalized to the ratio in non-infected cells (mean ± SEM, *n* = 3; *N* = 3). **M)** Aera under the curve of L (mean ± SD, *N* = 3, One-way ANOVA).

**Figure 8.**
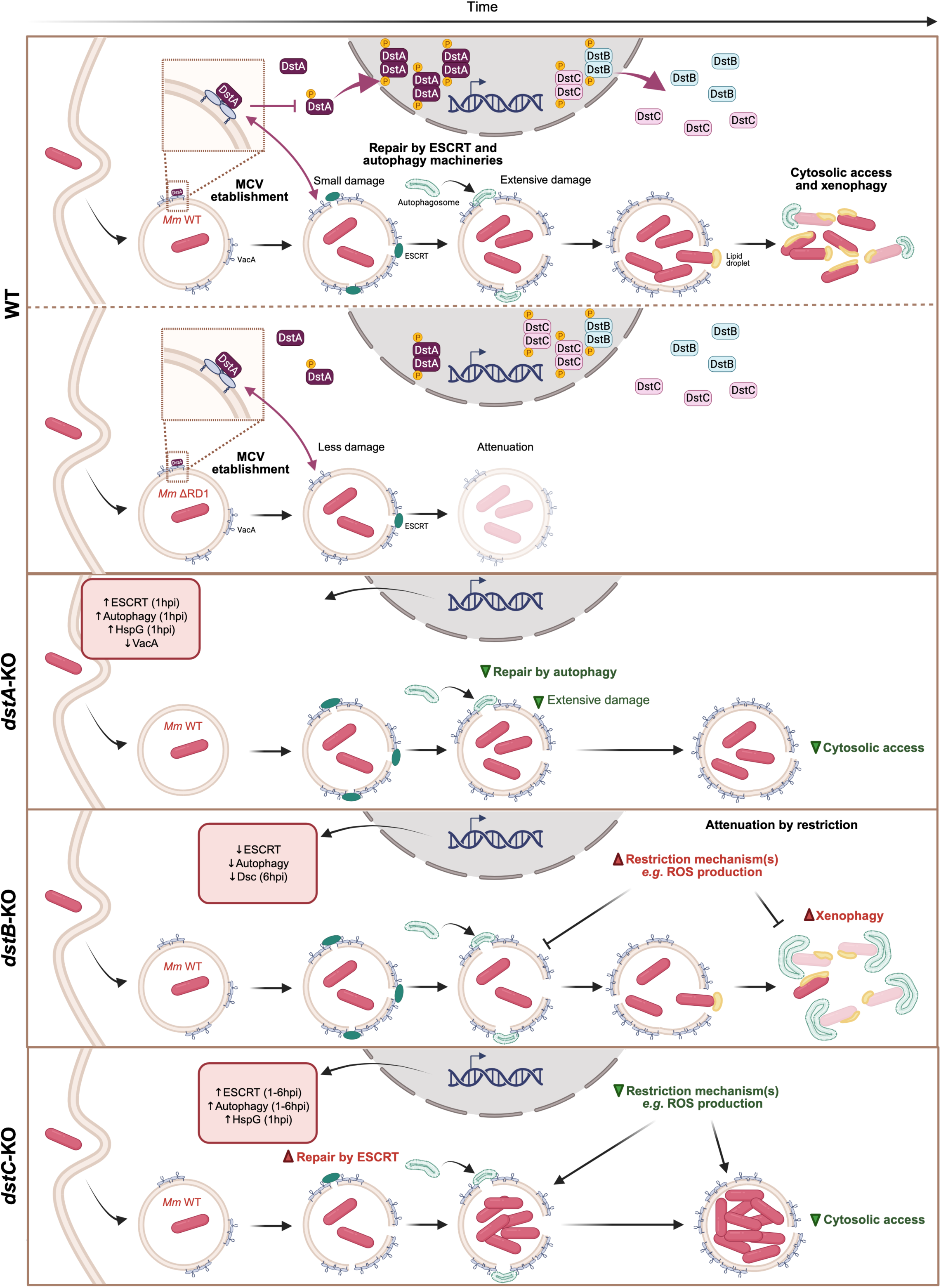
Working model of DstA, DstB and DstC-dependent regulation of MCV integrity and intracellular *Mm* fate. Schematic model illustrating the translocation dynamics of DstA, DstB and DstC during infection and the consequences of their absence. Following phagocytosis of *Mm*, DstA is rapidly and transiently translocated to the nucleus, whereas DstB and DstC progressively relocalize to the cytosol. At the early stage of infection, DstA interacts with VacA and then regulates its accumulation at the MCV during infection with both *Mm* WT and the damage-deficient mutant (ΔRD1). Conversely, VacA limits the nuclear translocation of DstA by acting as a cytoplasmic anchor. In *dstA*-KO cells, *Mm* induces less extensive damage to the MCV membrane, which subsequently leads to a reduced autophagy-mediated repair activity and its confinement within the MCV. In *dstB*-KO cells, *Mm* is restricted in the cytosol through an enhanced xenophagy restriction and increased ROS production. In *dstC*-KO cells, the ESCRT-mediated repair is more active, leading to the confinement of *Mm* within the MCV, however, *Mm* exhibits improved fitness in this compartment, due to a diminished ROS activity.

We next assessed whether reduced VacA expression affects its localization at the MCV. To this end, we quantified both the percentage of cells with GFP-VacA patches colocalizing with mCherry-expressing *Mm* and also measured the area of the GFP-VacA structures. In *dstA*-KO cells, the proportion of *Mm* WT-containing VacA-positive vacuoles was higher at 6 and 24 hpi compared to infected WT cells, although the size of VacA patches remained unchanged (Fig. 7f-h). Thus, beyond the positive transcriptional regulation of VacA, DstA appears to regulate the accumulation of VacA at the MCV. Surprisingly, *Mm* ΔRD1-containing vacuoles were more frequently coated with GFP-VacA, similar to *Mm* WT-containing vacuoles in WT cells. In addition, the VacA patch size on *Mm* ΔRD1-containing vacuoles was comparable to that observed in WT or *dstA*-KO cells infected with *Mm* WT (Fig. 7f-h). Previous studies showed that knockout of *vacuolins* reduces EsxA-mediated MCV damage and bacteria escape to the cytosol ^42^. In line with this, we observed that in *dstA*-KO cells, *Mm* remains confined within the MCV. Taken together, these findings and the increased frequency of VacA accumulation at the MCV in *dstA*-KO cells suggest that *Mm*-induced membrane damage activity is dampened by a disrupted balance of VacA-associated events at the MCV. Overall, our results indicate that altered VacA accumulation at the MCV membrane compromises EsxA-mediated damage activity, underscoring the importance of maintaining appropriate levels of VacA localization at this compartment. This study further highlights that DstA regulates VacA accumulation at the MCV, likely through direct interaction.

### VacA controls the early nuclear translocation of DstA

To further evaluate the functional interplay between DstA and VacA, we next examined whether VacA modulates the DstA nucleo-cytoplasmic distribution. For this purpose, we used *vacA*, *vacB* and *vacC* (*vacABC*) triple KO cells and co-expressed GFP-DstA and mCherry-H2B to quantify DstA nuclear translocation upon infection. The percentage of infected cells displaying active DstA nuclear translocation was higher after 2 hpi in *vacABC*-KO cells compared to *dstA*-KO cells expressing GFP-DstA (Fig. 7j-k). In addition, nuclear localization was amplified in *vacABC*-KO cells, resulting in more sustained nuclear translocation response in this background (Fig.7l-m). These results demonstrate that VacA negatively regulates DstA nuclear translocation, probably functioning as a cytoplasmic anchor to prevent its translocation in the nucleus. This also supports the existence of a regulatory loop in which DstA promotes VacA transcription and physically controls its accumulation at the MCV, while VacA, in turn, limits DstA nuclear translocation.

## Discussion

The absence of homologues of cytokine receptors, JAK kinases, and interferon cytokines, together with the conservation of downstream STAT signalling in *Dd*, prompted us to investigate the ancestral role of STAT proteins in antimycobacterial defence. We found that infection with *Mm* induces dynamic relocalization of DstA, DstB and DstC at distinct stages of infection. DstA rapidly translocates to the nucleus in a manner strictly dependent on *Mm*-induced membrane damage. In contrast, DstB and DstC undergo a slower redistribution to the cytosol, partially driven by bacteria-induced damage. Interestingly, the latter cytosolic redistribution is also observed in bystander cells, consistent with findings in mammals, where early STAT3 phosphorylation and nuclear translocation occur in bystander macrophages during *Mtb* infection ^14^. These observations suggest that DstB- and DstC-dependent signalling propagates via cell-to-cell communication, even in the absence of interferon secretion. This raises the question of how such communication operates within *Dd* populations, potentially through non-cytokine mediators such as damage-associated signals, extracellular vesicles, or direct cell-cell contact. One possibility is that mycobacterial cell wall components serve as signalling triggers. Indeed, *Mm* cell wall material actively traffics within infected cells and disseminates to bystander cells, priming them to resist subsequent infection ^61^. In a similar manner, release of *Mtb* cell wall fragments can prime resting neutrophils ^62^.

The ability of *Mm* to manipulate host defence mechanisms at early stages of infection is a decisive determinant of infection outcome, but the coordination of the underlying pathways is still unclear. Here, we demonstrate that DstA, DstB and DstC primarily function as transcriptional activators during early infection stages, each regulating distinct pathways, consistent with functional specialization in host defence. At early time points*, dstB-* and *dstC*-KO cells exhibit broad transcriptional downregulation of genes involved in transcription and protein synthesis, indicative of extensive host cell reprogramming. This response may potentially be linked to the complex regulation of the mTOR pathway ^32^, as well as to the documented lengthening of the cell cycle in both infected and bystander cells ^61^. At later stages, *dstB*-KO cells display altered expression of genes linked predominantly to central metabolism and membrane dynamics, whereas *dstC*-KO cells show defects in vitamin and lipid metabolism, as well as in membrane-associated processes. The regulation of genes associated with membrane dynamics likely reflects host responses to early infection events, including phagocytosis, MCV remodelling, lysosome fusion, xenophagy recapture of cytosolic bacteria, or early *Mm* exocytosis^63^. Collectively, these transcriptional analyses suggest that DstB and DstC play central roles in host transcriptional response to *Mm* invasion, thereby promoting metabolic adaptation and/or reinforcing antimicrobial defences. The identified interaction partners further point to specific stages of infection that may be regulated by these proteins. For instance, DstB may contribute to lectin-mediated cytosolic bacteria recognition. Concretely, DstB associates at early stages of infection with DscA, a cytosolic H-type lectin functionally homologous to galectins, which participates in the recognition of cytosolically exposed *Mm* glycolipids ^60^. Consistent with this observation, DstB also promotes the transcriptional upregulation of discoidin family members at 1 and 6 hpi. DscA was additionally found to interact with PefA under sterile damage induction. PefA is an ALG-2-like Ca^2+^ sensor proposed to orchestrate the recruitment of the E3 ubiquitin ligase TrafE, ESCRT components and the autophagy machinery to damaged membranes ^38^. Together, these interactions support the existence of an organized signalling network coordinating responses to membrane damage and restricting cytosolic *Mm*. The identification of two mitochondrial carrier family proteins as DstA interactors further suggests that DstA may be linked to mitochondrial transport or metabolic homeostasis. Given the central role of mitochondrial carriers in membrane energetics and metabolite exchange, DstA could participate in stress-adaptive mitochondrial functions during infection.

Intriguingly, members of the HspG family, which belong to the small heat shock proteins (sHSPs) family, were strongly downregulated during the early stages of infection in WT cells. A similar transcriptional signature was previously reported in *Dd* cells exposed to *Mm,* where 11 out of the 20 most downregulated genes encode small Hsp20-family proteins, including HspG family members ^64^. Comparable responses have also been observed in AGS gastric adenocarcinoma cells exposed to *Helicobacter pylori* ^65^. In our context, *dstA-* and *dstC*-KO cells displayed only a modest downregulation of these genes at 1 hpi, further reinforcing an important contribution of these proteins during the early stages of *Mm* infection. HspG proteins are molecular chaperones that maintain proteostasis under stress conditions, primarily by preventing protein aggregation and sequestering misfolded proteins ^52,55^. In addition, sHSPs participate in anti-oxidant and anti-apoptotic functions, and help preserve cytoskeletal integrity ^52–55^. For example, HSPB1 protects against oxidative stress by reducing ROS levels, promoting proteasome-dependent degradation of oxidized proteins, and protecting the cytoskeleton from oxidative damage ^52^. The altered regulation of sHSPs observed in *dstA*- and *dstC*-KO cells therefore suggests that these proteins may contribute to the control of oxidative stress and/or apoptotic responses during infection. Interestingly, the human autophagy adaptor SQSTM1 (also known as p62) recruits HSP27 to promote efficient lysophagy ^66^. In this pathway, lysosomal stress increases intracellular ROS levels, leading to activation of p38 mitogen-activated protein kinase (MAPK) and its downstream effector kinase MAP kinase-activated protein kinase 2 (MK2), which in turn phosphorylates HSP27 and promotes the engulfment and clearance of damaged lysosomes. A related molecular mechanism might operate during mycobacterial infection and may be particularly relevant in *dstA*-KO cells, where the weaker downregulation of HspG expression might enhance lysophagy, thereby contributing to the observed restriction of bacteria within the MCV and to the reduced intracellular bacteria growth. HSP27 has also been implicated in p38MAPK-dependent cytoskeletal reorganization during pemphigus vulgaris IgG-induced acantholysis ^67^, further supporting a potential broader role for sHSPs in antimycobacterial responses. In addition to their cytoprotective functions, several mammalian sHSPs associate with cellular membranes and have been proposed to stabilize membrane integrity under stress conditions ^68^. The role of sHSPs in maintaining membrane bilayer stability through specific lipid interactions was first demonstrated in *Synechocystis* PCC 6803 ^69^. These observations may also be linked to the altered membrane integrity observed in *dstA*-KO cells. Together, these findings highlight potentially important roles for sHSPs in the cellular responses to mycobacterial infection.

DstA, DstB and DstC were found to associate with four Gα subunits of the heterotrimeric G proteins signalling pathways, with DstA additionally interacting with a distinct family member. These findings suggest that Dst proteins are functionally linked to G protein signalling during early stages of infection. In *Dd*, Gα subunits function as central switches that translate extracellular cues, such as cAMP and folate, into cellular responses including chemotaxis, signal relay, gene expression and cell differentiation ^70–73^. Although the precise role of G protein signalling in the context of mycobacterial infection remains to be elucidated, these interactions point to a potentially important contribution during infection. Consistent with this hypothesis, we observed a specific transcriptional signature associated with these Gα subunits during infection. In agreement with a role for G protein signalling in host-pathogen interactions, loss of Gβ subunit, which broadly impairs Gγ signalling, causes defective uptake of *Klebsiella aerogenes* ^74^. In this context, bacterial surface ligands activate the G-protein-coupled receptor (GPCR) folic acid receptor 1 (fAR1), which recognizes both folate and lipopolysaccharide and signals through Gβγ and Gα subunits such as Gα4 (GpaD) to promote bacterial engulfment ^74,75^. Notably, GpaD, a common interactor of all three Dst proteins, displayed an opposing transcriptional profile and is upregulated during infection with damage-deficient *Mm* compared to infection with *Mm* WT, suggesting that its regulation depends on *Mm*-induced membrane damage. Since loss of GpaD impairs phagocytic uptake ^76^, these findings identify G protein signalling, and GpaD in particular, as a promising pathway for further investigation in the context of mycobacteria infection. More specifically, elucidating the role of phagosomal Gα signalling could provide important insights into host–pathogen interactions and the regulation of phagosome function during infection.

DstA regulates lysosomal processes at the transcriptional level and subsequently modulates fatty acid biosynthesis. In mammals, STAT proteins have only been marginally linked to lysosomal function. Notably, STAT1-deficient macrophages fail to properly acidify phagolysosomes during *Leishmania major* infection ^77^, and STAT6 can reprogram lysosomal gene expression in response to specific cytokines and pathogens ^78^. However, direct evidence for STAT-mediated regulation of phagosome-lysosome function during infection remains limited, underscoring an understudied potential conserved role for DstA in lysosomal control. In this study, we identified DstA as a interactor of Ap3d1, a component of the AP-3 adaptor complex involved in trafficking between trans-Golgi network, endosomes and lysosomes or lysosome-related organelles ^57^. DstA also interacts with PatA, a Ca^2+^-induced plasma membrane ATPase ^58^, further supporting a role for DstA in lysosomal regulation. Previously, we showed that *Mm* remodels the composition and properties of the MCV membrane through manipulation of host membrane microdomains proteins and lipids ^42^. This remodelling enhances EsxA-mediated damage and promotes bacterial escape from the vacuole. Our findings now identify an additional host regulatory mechanism operating at this phagosomal stage. DstA directly associates with VacA and transcriptionally upregulates *vacA* expression, thereby increasing VacA protein levels, and somehow promoting VacA accumulation at the MCV membrane. This interaction contributes to the control of MCV membrane integrity and modulates damage induced by *Mm*. This additional regulatory layer involves protein-protein interaction and a regulatory feedback loop, whereby VacA restricts DstA nuclear translocation, thereby limiting its own transcriptional upregulation. Notably, among the three vacuolin isoforms, DstA selectively regulates *vacA* expression without affecting *vacB* and *vacC* transcript levels. Despite the absence of isoform-specific enrichement at the MCV membrane ^42^, this selective regulation suggests a tightly orchestrated mechanism targeting a single component of the oligomeric complex. In the absence of DstA, VacA accumulates strongly at the MCV membrane in cells infected with damage-deficient *Mm*, indicating that DstA contributes to host discrimination between virulent and avirulent intracellular pathogens. Furthermore, DstA interacts with the ergosterol isomerase Erg2, raising the possibility that DstA-mediated changes in sterol metabolism additionally influence membrane organization and integrity, potentially through membrane microdomain remodelling. Together with VacA regulation, this mechanism may enable dynamic remodelling of membrane composition during infection. In summary, the present study identifies a regulatory mechanism underlying this process and uncover additional antimycobacterial processes coordinated by Dst proteins beyond their transcriptional activities, likely involving non-canonical activities. Collectively, our findings reveal ancestral regulatory roles of STAT proteins in host defences and provide new insight into the evolutionary origins of STAT-mediated responses during mycobacterial infections.

## Methods

### D. discoideum culture

All cell lines, listed in Supplementary Table 1, were axenically cultured in HL5c medium (Formedium) supplemented with penicillin (100 U/mL; Thermo Fisher) and streptomycin (100 μg/mL; Invitrogen), and maintained at 22°C. The *dst*-knock-out (KO) cell lines were generated in Ax2 (Ka) background ^79^ using CRISPR/Cas9 technology. The pTM1285-guide1-dstA plasmid was used to express a guide RNA (gRNA) targeting the exon 2 of *dstA* (TGATCACTTTGGTTTAATGG), pTM1285-guide2-dstB targets the exon 1 of *dstB* (TATCGGGTATAGATGATGAT), pTM1285-guide1-dstC targets the exon 2 of *dstC* (CAGCAACTCCACCAGCAATT) and pTM1285-guide1-dstD and pTM1285-guide2-dstD target the exon 1 of *dstD* (TGGACTAAACAGAATACAGT and CCAGTATTAGTTACAATCAT respectively). *Dd* were transfected with plasmids (Supplementary Table 2) by electroporation and selected with the relevant antibiotic. Hygromycin (InvivoGen) was used at a concentration of 50 μg/mL for cell lines expressing reporters at the safe-haven act5 locus. Blasticidin was used at a concentration of 5 μg/mL for VacA-GFP KI cell lines and mCherry-H2B and at act6 locus.

### Mycobacteria culture

*M. marinum* WT and ΔRD1 mutant were grown in Middlebrook 7H9 (Difco) supplemented with 10% Oleic Albumin Dextrose Catalase (OADC; Becton Dickinson), 0.2% glycerol, and 0.05% tyloxapol (Sigma Aldrich) at 32°C in shaking culture at 150 rpm in the presence of 5-mm glass beads to prevent clumping. Hygromycin was used at a concentration of 50 μg/mL for mCherry- or GFP-expressing *Mm* strains and at 100 μg/mL for *Mm* pTEC18. Kanamycin was used at 50 μg/mL for GFP- and lux-expressing *Mm* strains.

### *Dd* infection with *Mm*

Infections were performed as previously described ^25,80^. Briefly, *Mm* strains were cultured for 24 h prior to the experiment at 32°C under shaking conditions in 7H9 broth (Becton Dickinson, Difco Middlebrook 7H9) supplemented with 0.2% glycerol (Sigma-Aldrich), 10% OADC (Becton Dickinson), and 0.05% tyloxapol (Sigma-Aldrich). After bacterial growth, the OD_600_ was measured and adjusted to infect *Dd* cells, which had been plated the day before, at a multiplicity of infection (MOI) of 10. Following infection by spinoculation, extracellular bacteria were removed by washing, and adherent infected cells were resuspended in filtered HL5c medium supplemented with a bacteriostatic concentration of streptomycin (5 μg/mL) and penicillin (5 U/mL) to prevent extracellular *Mm* growth. Mock non-infected cells were treated identically but without bacteria. MβCD (2 mM; Sigma-Aldrich) was added immediately prior to infection. Intracellular bacterial growth was monitored by measuring the luminescence using a Synergy Mx Monochromator-Based Multi-Mode Microplate Reader (Biotek) at 25°C in a white 96-well plates.

### Western blotting

For certain western blot analyses, cells infected with GFP-expressing *Mm* were sorted by FACS prior to lysis. Collected cells were lysed in 1X Laemmli loading buffer (62.5 mM Tris pH 6.8, 2% (w/v) SDS, 10% glycerol, 0.005% (w/v) bromophenol blue, 5 mM DTT). For western blot analyses using a phospho-specific antibody, cells were washed with 1X Sorensen buffer (15 mM KH2PO4, 2mM Na2HPO4, 50 μM MgCl2, 50 μM CaCl2, pH 6.0) supplemented with 120 mM Sorbitol, 1X cOmplete^TM^ Inhibitor cocktail (Roche) and 1X PhosSTOPTM (Roche) prior to lysis in Laemmli loading buffer. Approximately 5x10^5^ cells corresponding to 25 μg of proteins were denatured 5 min at 95°C and SDS–polyacrylamide gel electrophoresis separation and transfer onto nitrocellulose membranes (Amersham Protran). Membranes were blocked during 1 hour at room temperature in 1X TBS containing 5% (w/v) BSA (SERVA Electrophoresis GmbH) or containing 5% (w/v) milk (GE Healthcare). Immunodetection was performed using the antibodies described in Supplementary Table 3 and the signal was revealed by enhanced chemiluminescence (ECL; Amersham Biosciences) using a Fusion Fx device (Vilbert Lourmat).

### Microscopy

For quantification of the GFP-Vps32, mCherry-Atg8a and mCherry-Plin reporters, 5 x 10^5^ cells were plated in 96-well IBIDI dishes in filtered HL5c medium. Cells were imaged at the indicated time points using a 60x water-immersion objective on an ImageXpress Micro XL HC microscope. For each condition, three wells were imaged with nine imaged fields per well and three z-sections per field acquired at 1-μm intervals. For translocation assay, 5 x 10^5^ cells were plated in 96-well IBIDI dishes and imaged every hour for 12 hours with a 60x water-immersion objective on the ImageXpress Micro XL HC microscope. For each condition, three wells were imaged with nine image fields per well acquiring three z-sections per field acquired at 1 μm intervals. For LLOMe experiments, 5 x 10^5^ cells were plated in 96-well IBIDI dishes and imaged every 5 minutes for 2 hours using a 60x water immersion objective on the ImageXpress Micro XL HC microscope. For each condition, two or three wells were imaged with four image fields per well and three z-sections per field acquired at 1-μm intervals. An initial image was acquired for all conditions before addition of 4.5 mM of LLOMe, after which the localization of GFP-DstA, mCherry-H2B and GFP-Vps32 was monitored. For MβCD treatment, cells were incubated with 2 mM MβCD for 40 min prior to LLOMe addition. Single-cell analyses were performed using MetaExpress software to quantify dot formation, area, or fluorescence intensity, as previously described ^81^. Image-based analysis were used to quantify the percentage of dot-positive cells or the nucleo-cytoplasmic ratio.

For imaging immediately after phagocytosis, cells were plated a μ-Slide 8-well IBIDI chambers, and *Mm* were added directly under the microscope. Images were acquired at 25°C using a 63x glycerol-immersion objective mounted on a spinning-disc confocal microscope (Intelligent Imaging Innovations Marianas SDC mounted on an inverted Leica DMIRE2 microscope).

### Image analysis

To quantify the percentage of *Dd* cells positive for GFP-Vps32, mCherry-Atg8a and VacA-GFP at the MCV, *Mm* was segmented based on its fluorescent tag (mCherry or GFP) and cells were segmented using either the bright-field or fluorescence channel depending on the expression level. Reporter recruitment was detected by first identifying structures with signal intensities higher than the cytosol and subsequently assessing their colocalization with the *Mm* mask based on pixel overlap with the bacterial signal. Plin-bacteria colocalization was analysed using a similar segmentation procedure. In this case, the percentage of surface overlap between the Plin mask area and the expanded bacterial mask area was quantified, with intracellular bacterial masks enlarged by one pixel. A similar segmentation procedure without a bacterial mask was used to identify GFP-Vps32 dot formation following LLOMe treatment.

To quantify GFP intensity of Dst proteins, cells were segmented based on the bright-field image. Infected cells were identified by the presence of intracellular bacteria. Using mCherry-H2B marker, nuclei were segmented, and their areas enlarged to avoid signal contamination from the bright nuclear region. Nuclear and cytosolic GFP-Dst intensities were then measured, with the cytosolic signal calculated excluding the enlarged nuclear mask. A similar segmentation procedure without bacteria mask was used to determine the nucleo-cytoplasmic ratio of GFP-DstA following LLOMe treatment.

### RNA sequencing sample preparation and analysis

Following infection of Dd WT and *dst*-KO cells with GFP-expressing *Mm*, infected and mock non-infected cells were pelleted, resuspended in 500 μL HL5c and sorted by FACS (Beckman CoulterMoFlo Astrios). Gating was performed based on cell size (forward scatter) and granularity (side scatter). Infected (GFP-positive) and non-infected (GFP-negative) sub-populations were defined according to GFP intensity in the FITC channel. Typically, ∼5 x 10^5^ cells of each fraction were collected for RNA isolation. RNA from mock non-infected or infected cells was extracted at the indicated time points using the Direct-zol RNA MiniPrep kit (Zymo Research) following the manufacturer’s instructions. Quality of RNA libraries, sequencing, and bioinformatic analyses were performed as previously described ^43^.

### GFP-trap

Approximately 3x10^7^ *Dd dst*-KO cells expressing GFP-DstA, GFP-DstB or GFP-DstC and infected with *Mm* mCherry were pelleted at 1 hpi and washed with Sorensen buffer supplemented with 120 mM of Sorbitol. Cells were lysed on ice using RIPA buffer (50 mM Tris-HCl pH 7.5, 150 mM NaCl, 0.1% SDS, 2 mM EDTA pH 8.0, 0.5% sodium deoxycholate, 0.5% Triton X-100) supplemented with protease and phosphatase inhibitor cocktails (Roche). Lysates were incubated at 4°C on rotating wheel for 45 min, and the supernatants were incubated with 35 μL GFP-Trap® Magnetic Agarose (ChromoTek) for 4 h at 4°C on a wheel. Beads were captured using a magnetic rack and washed twice in Triton X-100-containing buffer (50 mM Tris-HCl pH 7.5, 150 mM NaCl, 50 mM sucrose, 5 mM EDTA pH 8.0, 0.3% Triton X-100, 1 mM DTT, 5 mM ATP, protease and phosphatase inhibitors), followed by three washes in the same buffer without Triton X-100. Three independent biological replicates were processed for each bait.

### Liquid chromatography-electrospray ionization-tandem mass spectrometry

Samples were prepared using iST kits (Preomics) according to manufacturer’s instruction. Briefly, beads were re-suspended in 50 μL of provided lysis buffer and proteins were denatured, reduced and alkylated during 10 min at 60°C. The resulting slurries (beads and lysis buffer) were transferred to dedicated cartridges and proteins were digested with a Trypsin/LysC mix for 2 hours at 37°C. After two cartridge washes, peptides were eluted with 2 x 100 μL of provided elution buffer. Samples were finally completely dried under speed vacuum and stored at -20°C. Samples were dissolved in 20 μL of loading buffer (5% CH3CN, 0.1% FA) and 4 μL were injected on column. Liquid chromatography-electrospray ionization-tandem mass spectrometry (LC-ESI-MS/MS) was performed on a Q-Exactive HF Hybrid Quadrupole-Orbitrap Mass Spectrometer (Thermo Fisher Scientific) equipped with an Easy nLC 1000 liquid chromatography system (Thermo Fisher Scientific). Peptides were trapped on a Acclaim pepmap100, C18, 3μm, 75μm x 20mm nano trap-column (Thermo Fisher Scientific) and separated on a 75 μm x 250 mm, C18, 2μm, 100 Å Easy-Spray column (Thermo Fisher Scientific). The analytical separation was run for 90 min using a gradient of H2O/FA 99.9%/0.1% (solvent A) and CH3CN/FA 99.9%/0.1% (solvent B). The gradient was run as follows: 0-5 min 95 % A and 5 % B, then to 65 % A and 35 % B for 60 min, and 5 % A and 95 % B for 20 min at a flow rate of 250 nL/min. Full scan resolution was set to 60’000 at m/z 200 with an AGC target of 3 x 10^6^ and a maximum injection time of 60 ms. Mass range was set to 400-2000 m/z. For data dependent analysis, up to twenty precursor ions were isolated and fragmented by higher-energy collisional dissociation HCD at 27% NCE. Resolution for MS2 scans was set to 15’000 at m/z 200 with an AGC target of 1 x 10^5^ and a maximum injection time of 60 ms. Isolation width was set at 1.6 m/z. Full MS scans were acquired in profile mode whereas MS2 scans were acquired in centroid mode. Dynamic exclusion was set to 20s.

### Proteomic analysis

Raw data were processed using Proteome Discoverer 2.4 software (Thermo Fisher Scientific). Briefly, spectra were extracted and searched against the *Dictyostelium discoideum* reference database (Uniprot, 12’718 entries), the *Mycobacterium marinum* database (Uniprot, 5418 entries), the three known GFP-tagged bait protein sequences, and an in-house database of common contaminant using Mascot (Matrix Science, London, UK; version 2.6.2). Trypsin was selected as the enzyme, with one potential missed cleavage. Precursor ion tolerance was set to 10 ppm and fragment ion tolerance to 0.6 Da. Carbamidomethylation of cysteine was specified as fixed modification. Variable amino acid modification was oxidation (M), and phosphorylation (STY). Modification site probabilities were calculated with the IMP-ptmrs Node. Peptide-spectrum matches were validated using Percolator a target FDR of 0.01 and a Delta Cn of 0.5. For label-free quantification, features and chromatographic peaks were detected using the “Minora Feature Detector” Node with the default parameters. PSM and peptides were filtered with a false discovery rate (FDR) of 1%, and then grouped to proteins with again an FDR of 1% (strict) or 5% (relaxed) and using peptides with high confidence level. Low abundance resampling was used for missing value imputation. Both unique and razor peptides were used for quantitation and protein abundances are calculated as the average of the three most abundant distinct peptide group. The abundances were normalized on the “Total Peptide Amount” and then “Protein abundance based” option was selected for protein ratio calculation and associated *p*-values were calculated with an ANOVA test (individual proteins).

### Statistical analyses

The sample size and *P*-values are presented in appropriate figure legends (*n* = number of technical replicates; *N* = number of biological replicates). Data are presented as means and SD (standards deviations) or SEM (standard errors of the mean) and were analysed with GraphPad Prism 10 software.

## Data Availability

The RNA-seq raw data will be available in a SRA/GeO Bioproject, and the code used for DEG analyses as well as the processed data will be available at Zenodo. In addition, the code used for RNA-seq analyses is partly available via the UNIGE bioinformatics platform (https://unige.ch/medecine/bioinformatics/home).

## Acknowledgements

We thank the Geneva Antibody facility, the Photonic Bioimaging Center, and the iGE3 Genomics Platform at the Faculties of Sciences and Medicine of the University of Geneva. We gratefully acknowledge ACCESS Geneva, with a special thanks to Dr. Dimitri Moreau for assistance with high-content microscopy experiments. This work was supported by Swiss national Science Foundation grants 310030_188813 and 310030_219364.

## Author Contributions

J.T. and T.S. conceived the project. J.T and N.H. performed the experiments. J.T. and N.H. analysed the data. L.R. analysed the RNA-sequencing data, and L.V. performed *in silico* DNA-binding motif prediction analysis. J.T. and T.S. wrote the manuscript with input from all authors. J.T. and T.S. supervised the study.

## Materials & Correspondence

All materials generated in this study are available upon request.

## Supplementary Information

### Supplementary Figures

**Figure S1.**
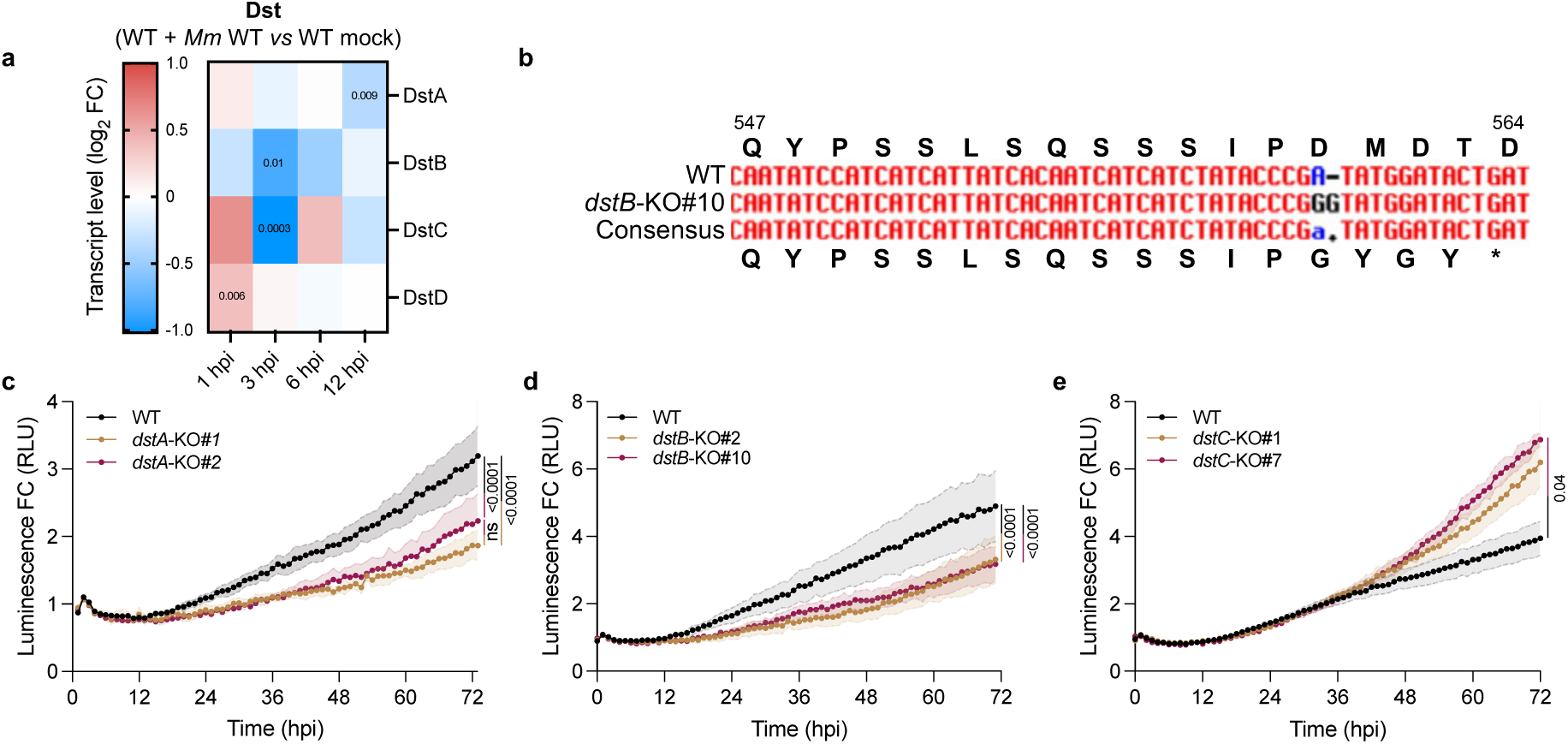
*Mm* intracellular growth is impacted in *dstA*-, *dstB*- and *dstC*-KO cells. **A)** Heatmap showing RNA-seq transcriptome analysis for Dst gene expression. RNA-seq data from infected *Dd* WT cells are normalized to non-infected WT cells and is represented as log_2_ fold change (N = 3). **B)** DNA sequencing of the genomic region encompassing the CRISPR-Cas9 target site. The corresponding protein sequence is shown above and below the DNA sequence. **C-E)** *Dd* WT, *dst*-KO and *dst*-KO cells expressing GFP-Dst were infected with bioluminescent *Mm* WT. Bioluminescence was monitored over 72 hours (mean fold change ± SEM, *n* = 3, *N* = 3, one-way ANOVA with Tukey’s multiple comparisons test). RLU, relative luminescence units.

**Figure S2.**
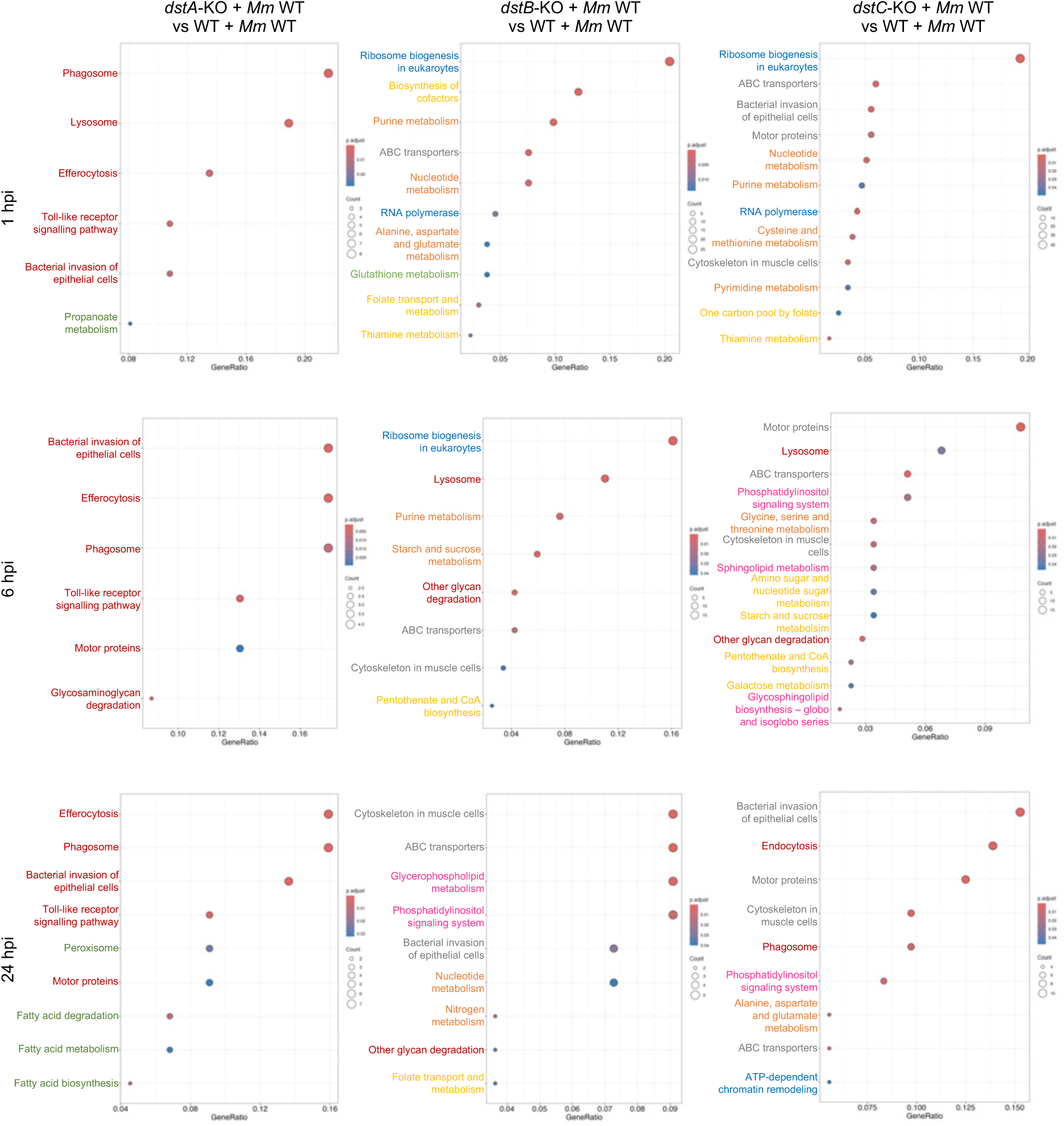
KEGG pathway enrichment analysis of downregulated genes compared to infected *Dd* WT. Significantly enriched pathways were identified using KEGG and ranked by adjusted *P* value. Dot size indicates the number of genes assigned to each pathway, and the x-axis denotes the gene ratio. Enriched pathway clusters are color-coded as follows: red, lysosomal/endocytic pathways; green, fatty acid metabolism; blue, gene expression and protein synthesis; orange, central metabolism and biosynthesis; yellow, cofactor and vitamin metabolism; fluorescent green, redox balance and stress response; grey, membrane transport and cytoskeleton; pink, lipid metabolism and membrane signalling.

**Figure S3:**
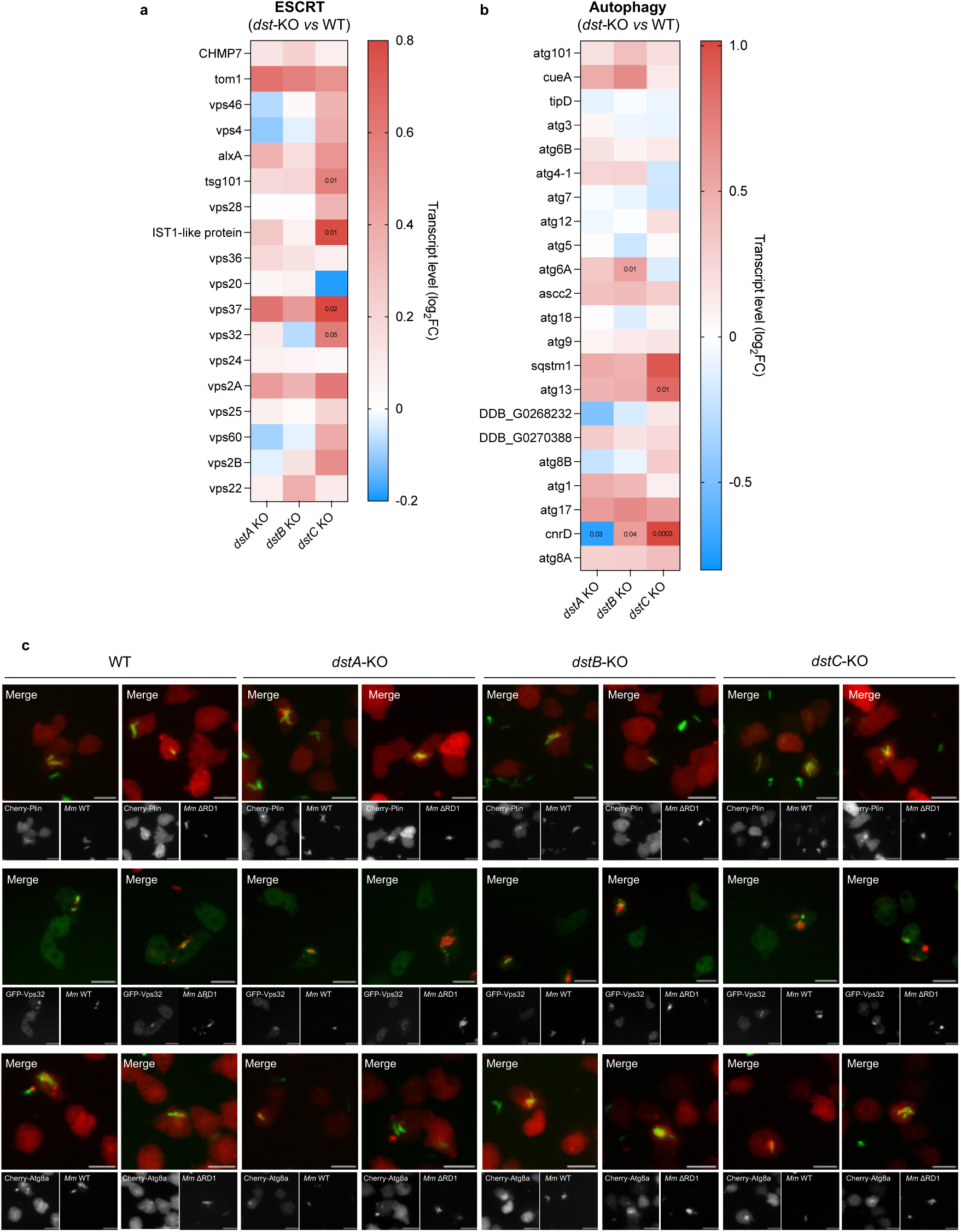
Transcriptional expression of autophagy and ESCRT-related genes at steady state and analysis of Plin, Vps32 and Atg8a reporters during infection in *dstA-, dstB- and dstC*-KO cells. **A-B)** Heatmaps showing RNA-seq transcriptome analysis for **A)** ESCRT- and **B)** autophagy-mediated repair gene expression at steady state. RNA-seq data from *Dd dst-*KO cells are normalized to *Dd* WT cells and is represented as log_2_ fold change (*N* = 3). **C)** *Dd* WT and *dst*-KO cells expressing mCherry-Plin, GFP-Vps32 or mCherry-Atg8a were infected with GFP- or Cherry-expressing *Mm* WT or ΔRD1 mutant and imaged live by high-content microscopy. Representative images are shown at 9 hpi for the Plin reporter, 1 hpi for the Vps32 reporter, and 24 hpi for the Atg8a reporter. Scale bars: 10 μm.

**Figure S4.**
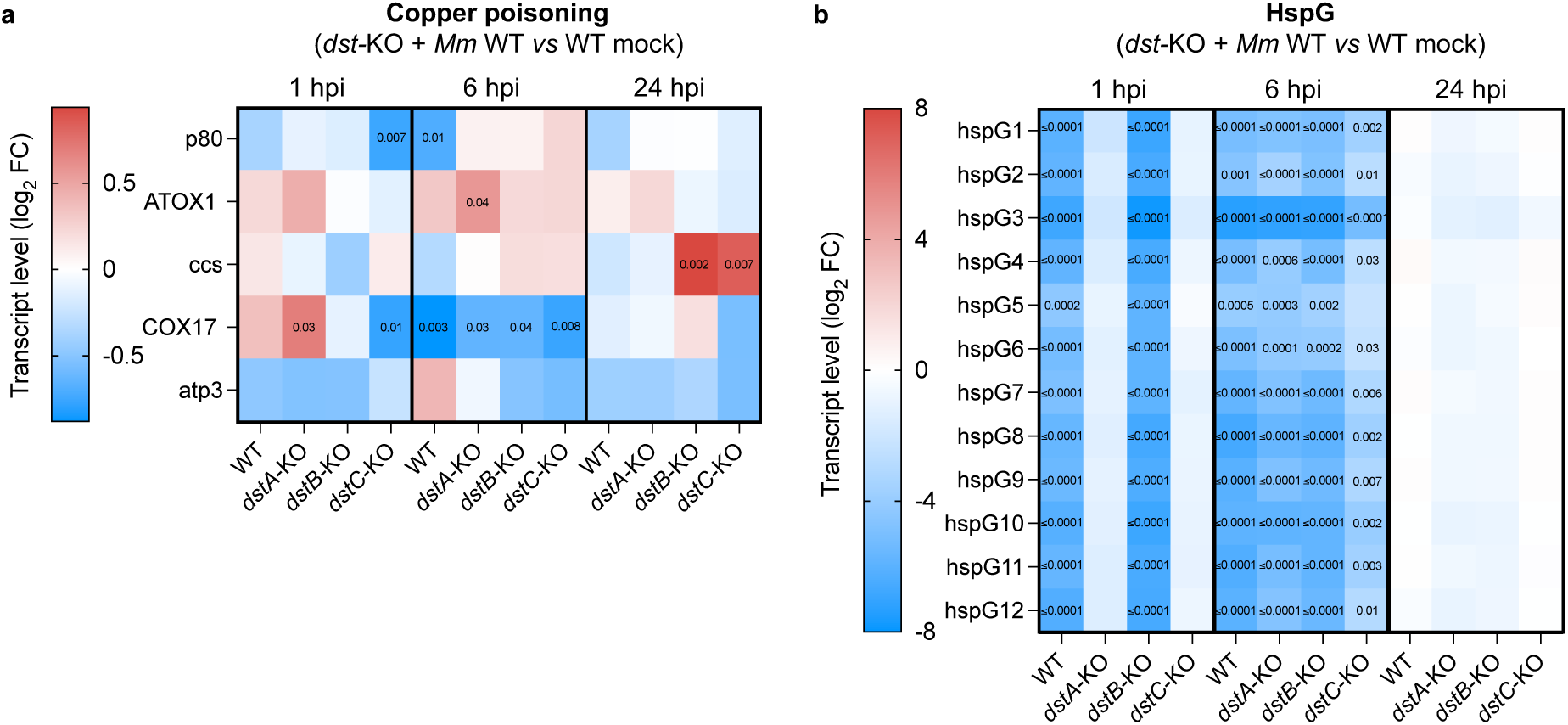
Transcriptional expression of copper poisoning- and HspG family-related genes in *dstA-*, *dstB-* and *dstC*-KO cells during infection. **A-B)** Heatmaps showing RNA-seq transcriptome analysis for **(A)** copper poisoning and **(B)** HspG family gene expression during infection. RNA-seq data from infected *Dd dst-*KO cells are normalized to non-infected *Dd* WT cells and is represented as log_2_ fold change (*N* = 3).

**Figure S5.**
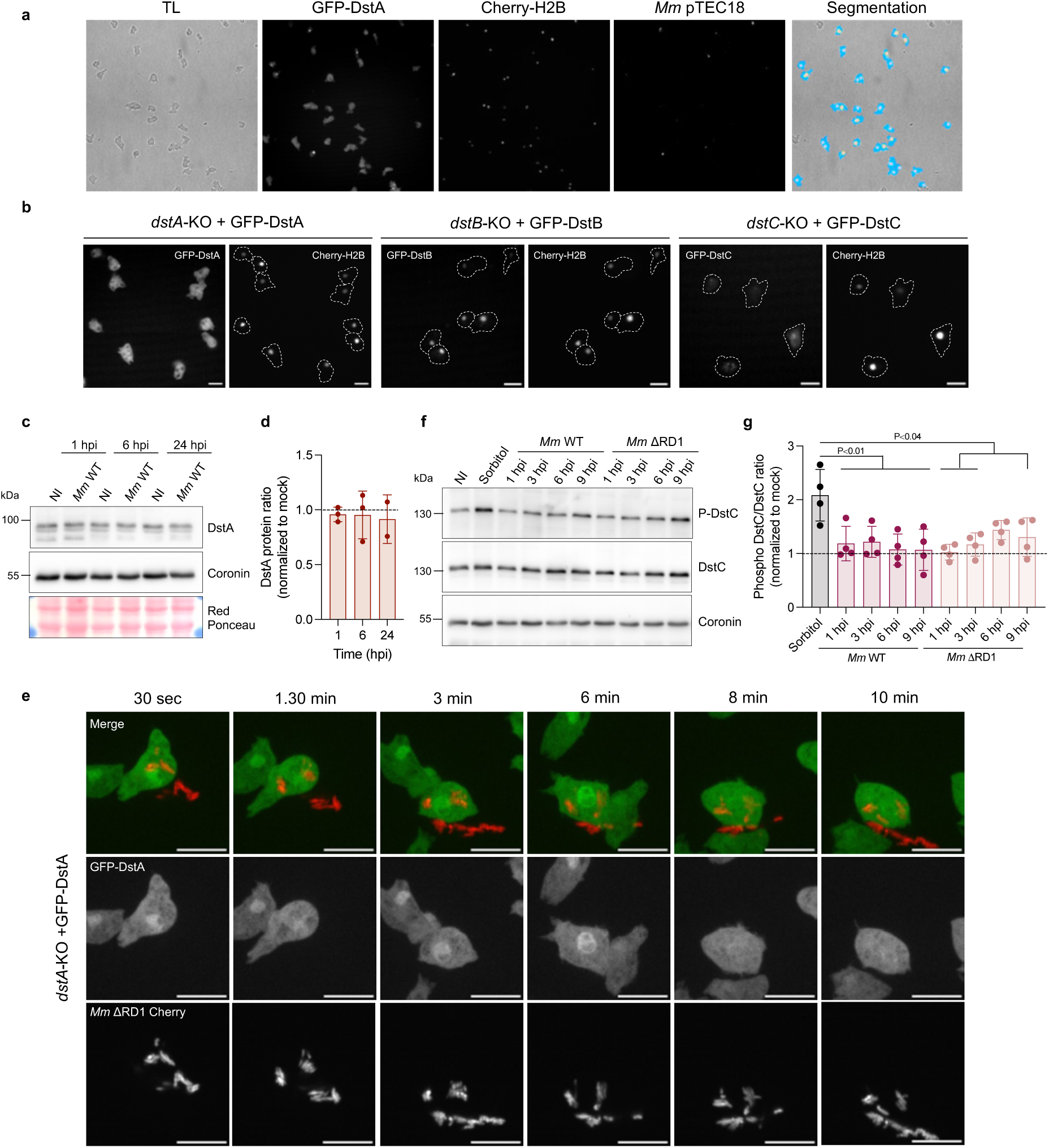
Membrane damage decreases DstC phosphorylation while DstA protein level is unchanged during infection. **A)** *Dd dstA*-KO cells expressing GFP-DstA and mCheny-H2B were infected with blue-fluorescent pTEC18 *Mm* WT and imaged by high-content microscopy. Representative live images at 1.5 hpi. Blue masks indicate the segmented cytosolic area excluding an enlarged nuclear mask (red + yellow/cyan) to avoid signal contamination from the bright nucleus. White masks identify the bacterium. Yellow masks are assigned to infected cells, whereas cyan masks to non-infected cells. GFP intensity in the nucleus and cytosol was used for quantification as a function of the infection status of each cell. **B)** *Dd ds*t-KO cells expressing GFP-Dst and mCheny-H2B were imaged at steady state by high-content microscopy. Scale bars: 10 μm. **C)** Representative immunoblot of *Dd* WT cells infected with WT *Mm* WT or under non-infected mock conditions, using the indicated antibodies. Infected cells were sorted by FACS prior to analysis. **D)** Quantification of DstA proteins represented in C) normalized to Coronin and mock condition (dashed line, mean ± SD, *N* = 3, one-way ANOVA). **E)** Live microscopy of *Dd dstA*-KO cells with mCheny-expressing *Mm* ΔRD1 mutant shortly after phagocytosis. Following incubation with *Mm*, infected cells was imaged for 10 min. Scale bars: 10 μm. **F)** Representative immunoblot of *Dd* WT cells infected with WT *Mm* WT or the ΔRD1 mutant, *Dd* WT treated with 120 mM of sorbitol as a positive control, or maintained under mock non-infected conditions, using the indicated antibodies. **G)** Quantification of phospho-DstC levels shown in F), normalized to total DstC protein and Coronin, and expressed relative to the mock condition (dashed line, mean ± SD, *N*= 4, one-way ANOVA).

**Figure S6.**
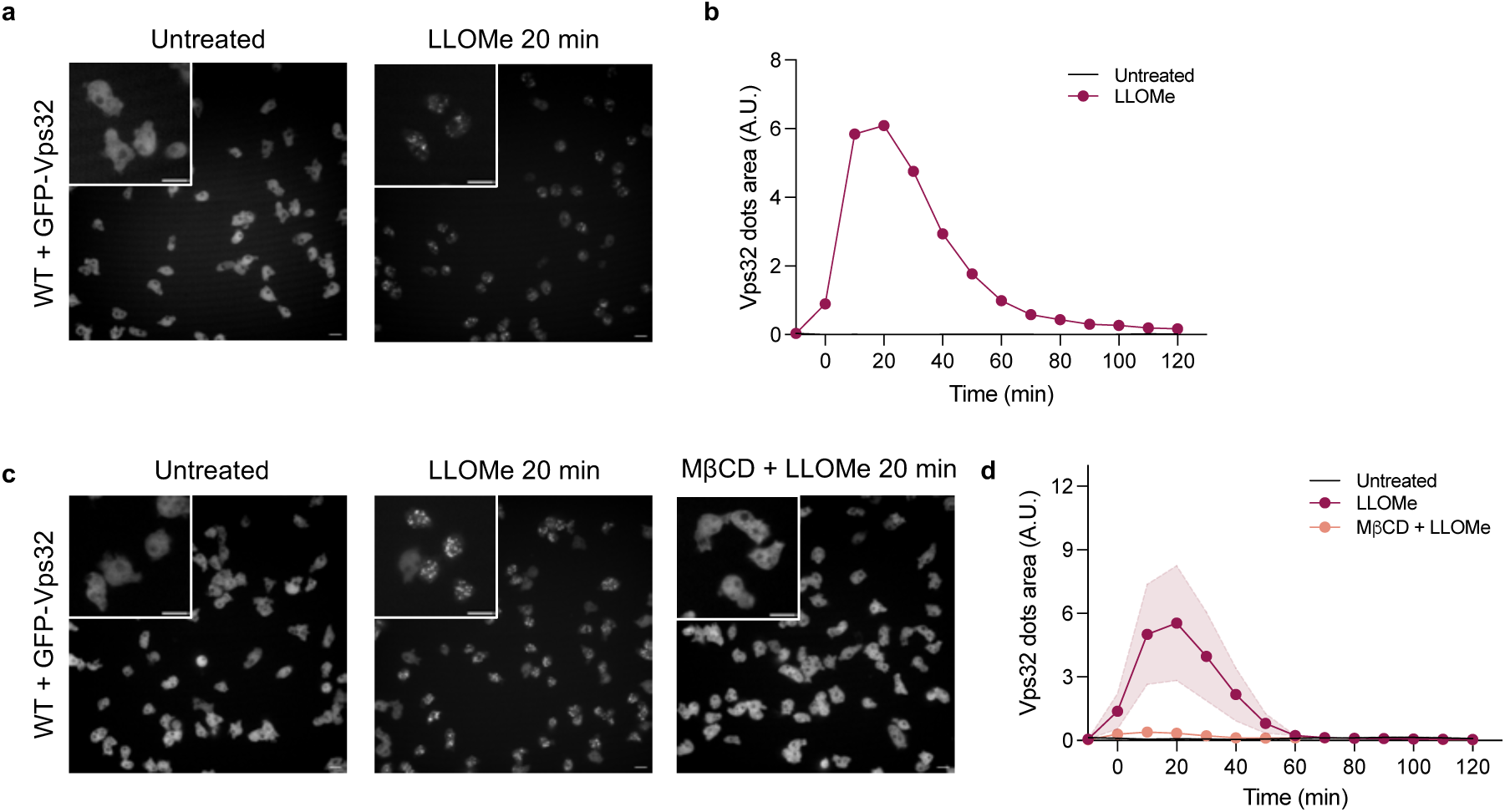
Recruitment of Vps32 after LLOMe treatment. **A-B)** *Dd* WT cells expressing GFP-Vps32 were treated with 4.5 mM of LLOMe or left untreated. **A)** Representative live images showing GFP-Vps32 localization before treatment and 20 min after LLOMe addition. Scale bars: 10 μm. **B)** Quantification of A. showing the area of GFP-Vps32 dots (mean ± SD. *N*= 1). **C-D)** *Dd* WT cells expressing GFP-Vps32 were pre-treated with 2 mM of MβCD and subsequently treated with 4.5 mM of LLOMe or left untreated. D) Quantification of C, showing the area of GFP-Vps32 dots (mean ± SD, *N* = 3)

**Figure S7.**
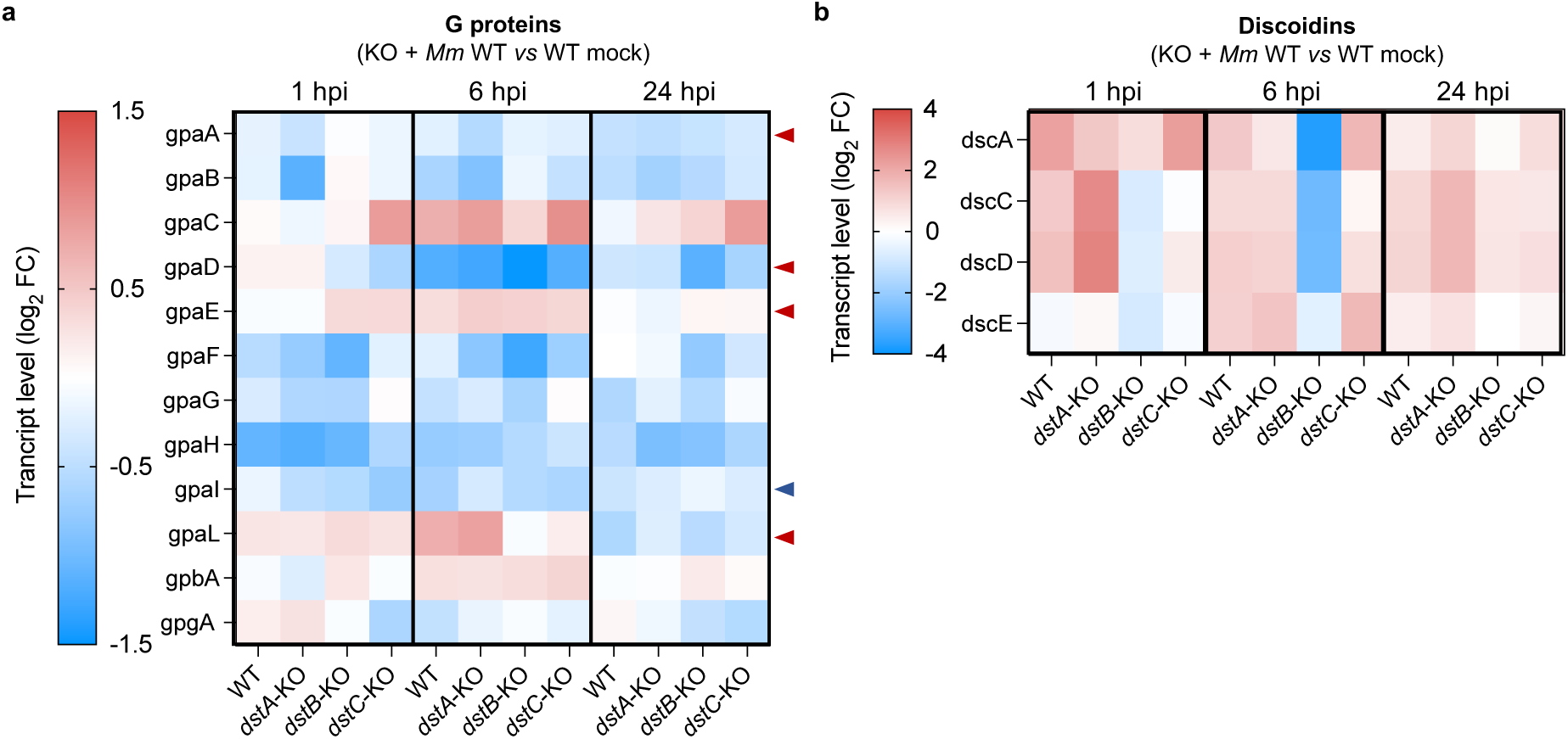
Transcriptional expression of G protein and discoidin genes. **A-B)** Heatmaps showing RNA-seq transcriptome analysis for **A)** *Gα* (*gpa*), Gβ (*gpbA*) and *Gγ* (*gpgA*) subunits and **B)** *discoidins* gene expression during infection. RNA-seq data from infected *Dd dst-*KO cells are normalized to non-infected *Dd* WT cells and is represented as log_2_ fold change (*N* = 3). Red arrows indicate Gα subunits that interact with DstA, DstB and DstC at 1 hpi. The blue arrow indicates Gα subunit that interacts with DstA.

**Figure S8.**
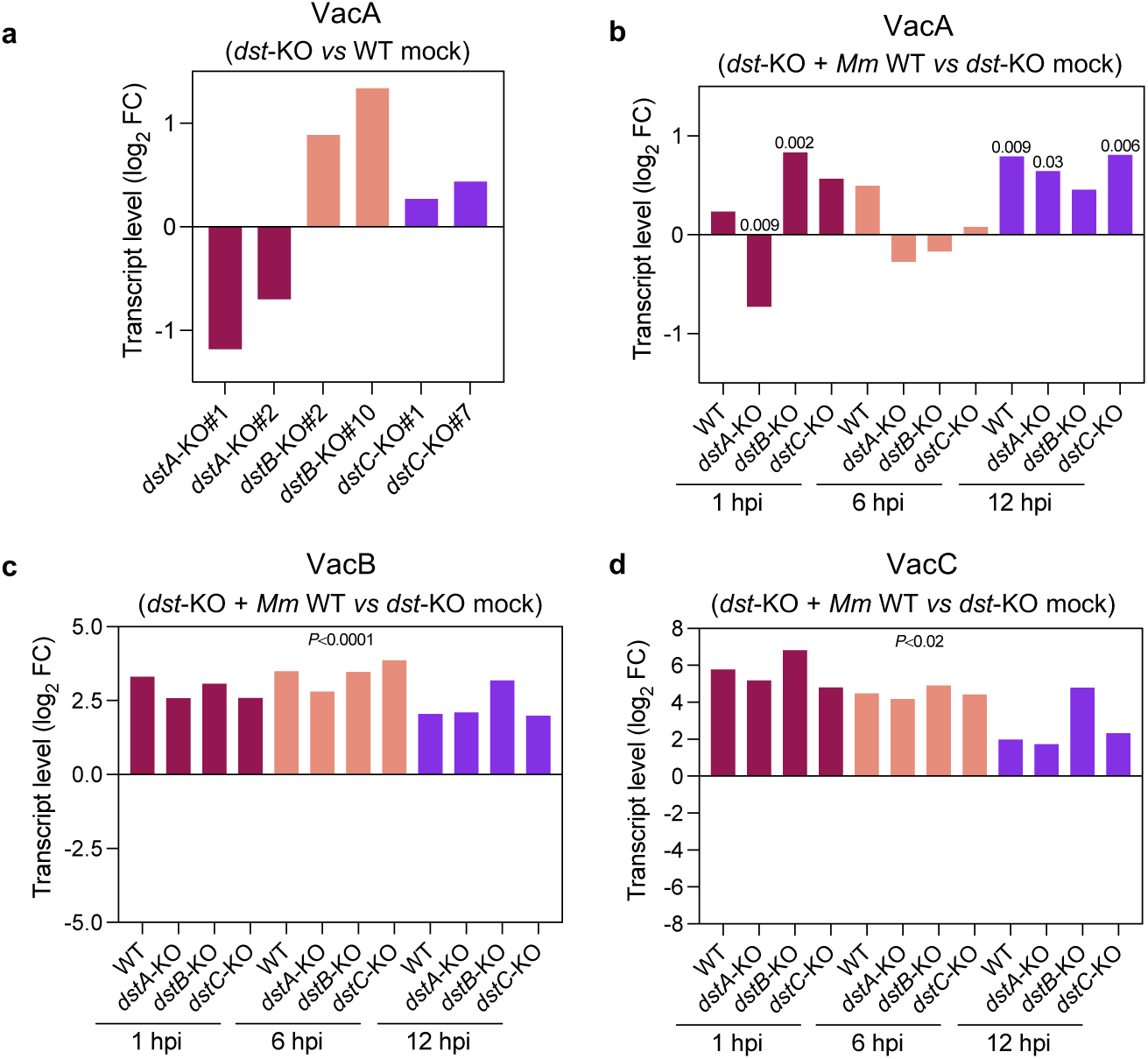
Transcriptional expression of *vacuolins* genes at steady state and during infection. **A)** RNA-seq analysis of *vacA* expression in *Dd dst*-KO cells under steady state condition, shown as fold changes relative to non-infected WT cells (mock, *N* = 3). **B-D)** RNA-seq analysis of *vacuolin* expression in FACS-sorted *Dd* WT or *dst*-KO cells infected with GFP-expressing *Mm* at the indicated hpi. Fold changes in **B)** *vacA*, **C)** *vacB* and **D)** *vacC* transcript levels are shown relative to the corresponding non-infected cells (mock, *N* = 3).

### Supplementary Tables

**Supplementary Table 1.** *D. discoideum* and *M. marinum* strains used in this study.

| Experimental models | Reference or source | Characteristic |
| --- | --- | --- |
| *Dd* Ax2 (Ka) WT |  |  |
| *Dd* Ax2 (Ka) *dstA*-KO#1 | This study |  |
| *Dd* Ax2 (Ka) *dstA*-KO#2 | This study |  |
| *Dd* Ax2 (Ka) *dstB*-KO#2 | This study |  |
| *Dd* Ax2 (Ka) *dstB*-KO#10 | This study |  |
| *Dd* Ax2 (Ka) *dstC*-KO#1 | This study |  |
| *Dd* Ax2 (Ka) *dstC*-KO#7 | This study |  |
| *Dd* Ax2 (Ka) *dstD*-KO#1 | This study |  |
| *Dd* Ax2 (Ka) *dstD*-KO#1 | This study |  |
| *Dd* Ax2 (Ka) *dstA*-KO + GFP-DstA | This study | act5::GFP-DstA |
| *Dd* Ax2 (Ka) *dstB*-KO + GFP-DstB | This study | act5::GFP-DstB |
| *Dd* Ax2 (Ka) *dstC*-KO + GFP-DstC | This study | act5::GFP-DstC |
| *Dd* Ax2 (Ka) *dstD*-KO + GFP-DstD | This study | act5::GFP-DstD |
| *Dd* Ax2 (Ka) *dstA*-KO + GFP-Vps32 | This study | act5::GFP-Vps32 |
| *Dd* Ax2 (Ka) *dstB*-KO + GFP-Vps32 | This study | act5::GFP-Vps32 |
| *Dd* Ax2 (Ka) *dstC*-KO + GFP-Vps32 | This study | act5::GFP-Vps32 |
| *Dd* Ax2 (Ka) *dstA*-KO + mCherry-Atg8a | This study | act5::mCherry-Atg8a |
| *Dd* Ax2 (Ka) *dstB*-KO + mCherry-Atg8a | This study | act5::mCherry-Atg8a |
| *Dd* Ax2 (Ka) *dstC*-KO + mCherry-Atg8a | This study | act5::mCherry-Atg8a |
| *Dd* Ax2 (Ka) *dstA*-KO + mCherry-Plin | This study | act5::mCherry-Plin |
| *Dd* Ax2 (Ka) *dstB*-KO + mCherry-Plin | This study | act5::mCherry-Plin |
| *Dd* Ax2 (Ka) *dstC*-KO + mCherry-Plin | This study | act5::mCherry-Plin |
| *Dd* Ax2 (Ka) *dstA*-KO + GFP-DstA + mCherry-H2B | This study | act5::GFP-DstA<br>act6::mCherry-H2B |
| *Dd* Ax2 (Ka) *dstB*-KO + GFP-DstB + mCherry-H2B | This study | act5::GFP-DstB<br>act6::mCherry-H2B |
| *Dd* Ax2 (Ka) *dstC*-KO + GFP-DstC + mCherry-H2B | This study | act5::GFP-DstC<br>act6::mCherry-H2B |
| *Dd* Ax2 (Ka) WT + GFP-DstA + mCherry-H2B | This study | act5::GFP-DstA<br>act6::mCherry-H2B |
| *Dd* Ax2 (Ka) WT + VacA-GFP KI | <sup>1</sup> |  |
| *Dd* Ax2 (Ka) *dstA*-KO + VacA-GFP KI | This study |  |
| *Mm* M strain |  |  |
| *Mm* ΔRD1 | | |

**Supplementary Table 2.** Plasmids used in this study.

| Plasmids | Reference or source | Characteristic |
| --- | --- | --- |
| pTM1285-guide1-dstA | This study | KO of *dstA* exon 2 |
| pTM1285-guide2-dstB | This study | KO of *dstB* exon 1 |
| pTM1285-guide1-dstC | This study | KO of *dstC* exon 2 |
| pTM1285-guide1-dstD | This study | KO of *dstD* exon 1 |
| pTM1285-guide2-dstD | This study | KO of *dstD* exon 1 |
| pDM1513-GFP-DstA | This study | act5::GFP-DstA |
| pDM1513-GFP-DstB | This study | act5::GFP-DstB |
| pDM1513-GFP-DstC | This study | act5::GFP-DstC |
| pDM1513-GFP-DstD | This study | act5::GFP-DstC |
| pDM1513-GFP-Vps32 | <sup>2</sup> | act5::GFP-Vps32 |
| pDM1514-mCherry-Atg8a | <sup>3</sup> | act5::mCherry-Atg8a |
| pDM1514-mCherry-Plin | <sup>4</sup> | act5::mCherry-Plin |
| pDMact6-Cherry-H2B | This study | act6::mCherry-H2B |

**Supplementary Table 3.** Antibodies used in this study.

| Antibodies | Reference or source | Catalog # |
| --- | --- | --- |
| Human Anti-DstA | <sup>5</sup> | RB865, synthesised at the Geneva Antibody Facility |
| Human Anti-DstC | <sup>6</sup> | RB874, synthesised at the Geneva Antibody Facility |
| Human Anti-DstD | <sup>7</sup> | RB878, synthesised at the Geneva Antibody Facility |
| Mouse Anti-Phospho-DstC CP22 | Jeffrey G. Williams lab | <sup>8</sup> |
| Mouse Anti-GFP | Abmart | M20004M |
| Mouse Anti-Coronin | Geneva Antibody Facility | ABCD_AK425 |
| Goat Anti-Mouse IgG (H + L)-HRP conjugate | Bio-rad | 1706516 |
| Goat Anti-Human (H+L) HRP conjugate | Bio-rad | 1721050 |

